# M4 drug discovery: human drug predictions from integrated preclinical insights exemplified with a GLP-1R agonist

**DOI:** 10.1101/2025.11.03.686224

**Authors:** Oscar Silfvergren, Sophie Rigal, Katharina Schimek, Christian Simonsson, Kajsa P Kanebratt, Felix Forschler, Burcak Yesildag, Uwe Marx, Liisa Vilén, Peter Gennemark, Gunnar Cedersund

## Abstract

A major recent breakthrough in the treatment of type 2 diabetes has been the development of glucagon-like peptide-1 receptor agonists (GLP-1RAs). However, current translational frameworks struggle to predict the clinical outcomes of these drugs from preclinical data. There are several reasons for this struggle, which are generic for many drugs: GLP-1RAs act through multi-timescale mechanisms in which short-term effects propagate into long-term changes; no single preclinical system can capture all their effects in humans; and mechanistic extrapolation requires modelling numerous whole-body biological processes. To address this gap, we present a new extrapolation approach, M4 drug discovery, and retrospectively apply it to the GLP-1RA exenatide in a manner that is generalisable to other drugs. The method integrates: Multi-level data (cellular to whole-body), Multi-timescale data (minutes to months), Multi-species data (e.g., rodents to humans), and Mechanistic knowledge. In this study, we integrate human cell and animal data with drug-free human studies to successfully predict human pharmacokinetics (cost < χ^2^, p=0.05; 64 < 97) and the outcomes of a 30-week clinical trial (36 < 45). We found that integrating information across the four M4 axes improved predictive performance and physiological relevance: multi-species data inform pharmacokinetics, human cell data provide human population- and donor-specific potency estimates, animal data reveal additional drug effects not observable in cell cultures, and the multi-timescale mathematical modelling enables short-term effects of exenatide and meals to inform long-term changes in insulin sensitivity. This work provides new tools for drug extrapolations, supporting the community towards safer and more informed preclinical-to-clinical drug extrapolations.

## INTRODUCTION

Diabetes and its associated pathologies constitute one of the most important disease areas, for which new drugs are urgently needed. A major recent breakthrough has been the development of glucagon-like peptide-1 receptor agonists (GLP-1RA), such as the GLP-1RA Ozempic (1,2). Today, many related therapies are developed, and their clinical utility is evaluated by predicting clinical outcomes from preclinical data. However, current translational frameworks struggle to make these extrapolations for GLP-1RAs (3–6). There are several reasons for this, which are applicable to many related therapies: they have multi-timescale mechanisms of actions (e.g., rapid insulin secretion and slow insulin sensitivity changes) that are difficult to mechanistically describe, as this requires stiff systems in which short-term effects accumulate into long-term changes; a mechanistic extrapolation of GLP-1RAs requires modelling many whole-body biological components (e.g., glucose–insulin interplay and appetite–diet changes); and no single preclinical system can capture all their effects in humans. These preclinical systems include conventional cell-based assays, microphysiological systems (MPS), and animal models, each with distinct strengths and limitations in mimicking human responses.

The strengths of cell-based assays are that they are inexpensive, fast, ethically viable, and provide a platform to study intracellular mechanisms. Unfortunately, traditional cell models lack 3D structure, organ–to-organ crosstalk, and a physiological context for pharmacokinetic properties. Conversely, MPS are more expensive, but they enable the study of interconnected *in vitro* organ models, thereby supporting investigation of human-like aetiologies and organ–organ crosstalk. We have previously developed a two-organ MPS containing 3D cell culture models of the liver and pancreas (7–10), and used this system to study the pharmacological effects of the GLP-1RA exenatide (11). We found that the MPS captures multi-timescale GLP-1RA-relevant mechanisms, including the rapid effect of exenatide on insulin secretion as well as the slow effect of diabetogenic conditions on insulin resistance. Together, both cell-based assays and MPS share an important strength: they enable the study of pharmacological effects and population variability using human cells prior to clinical testing. Finally, animal models often fail to accurately represent human responses and diseases, as seen, for example, in metabolic dysfunction models for rodents (12,13), but they provide a platform for studying pharmacology *in vivo* in a whole-body physiological context. In summary, cell-based assays, MPS, and animal systems each possess different strengths, and integrating their diverse features into a single framework could improve preclinical-to-clinical drug predictions. A powerful tool that potentially could integrate such diverse datasets into a single framework is mathematical modelling.

A computational model is a mathematical abstraction of a system (14–16), and these models are increasingly used in preclinical-to-clinical drug translations (17–19). One type of such mathematical model is ordinary differential equation (ODE) models, which have shown potential for describing human cell data and extrapolating it to humans, for example in drug-induced toxicity (20) and diarrhea (21). Both these studies use MPS to make human predictions, but lack the ability to consider organs not present in the MPS. There also exist ODE models that extrapolate whole-body pharmacokinetics from animals to humans (22,23), models that use both cell and animal measurements to predict human responses (19,24), and models that accumulate long-term changes from short-term dynamics (25). Yet, no model currently addresses all features simultaneously for preclinical-to-clinical predictions of GLP-1RAs and its effects on, for example, the glucose metabolism. We have previously developed multi-timescale glucose-oriented metabolism models for MPS (9,11), rodents (26), and humans (27,28). These models display three important features: a) <u>M</u>echanistic (describing the mechanistic underpinning of sampled data); b) <u>M</u>ulti-level (intracellular to whole-organism responses); and c) <u>M</u>ulti-timescale (seconds to months). These models can therefore be referred to as <u>M3</u>-models. Nonetheless, they are not yet connected along a translational axis, which hinders the leverage of complementary strengths across preclinical systems. Addressing this would require <u>M</u>ulti-species M3-models—that is, <u>M4</u>models.

We herein present a first M4 model of the GLP-1RA exenatide, and exemplify its utility in drug discovery by making clinical predictions from preclinical data (**Fig. 1**). Building on our foundational model of exenatide effects in human cells (11), we demonstrate how complementary preclinical insights, from human cells systems and animals studies, can be combined with mathematical modelling of drug-free human data to inform long-term effects from short-term drug responses. In doing so, we address complexities that current translational frameworks for GLP-1RAs struggle to handle (3–6). This broadly applicable approach strengthens the interpretation and translation of diverse preclinical systems and datasets, thereby informing decisions about advancing candidates into clinical studies.

**Figure 1.**
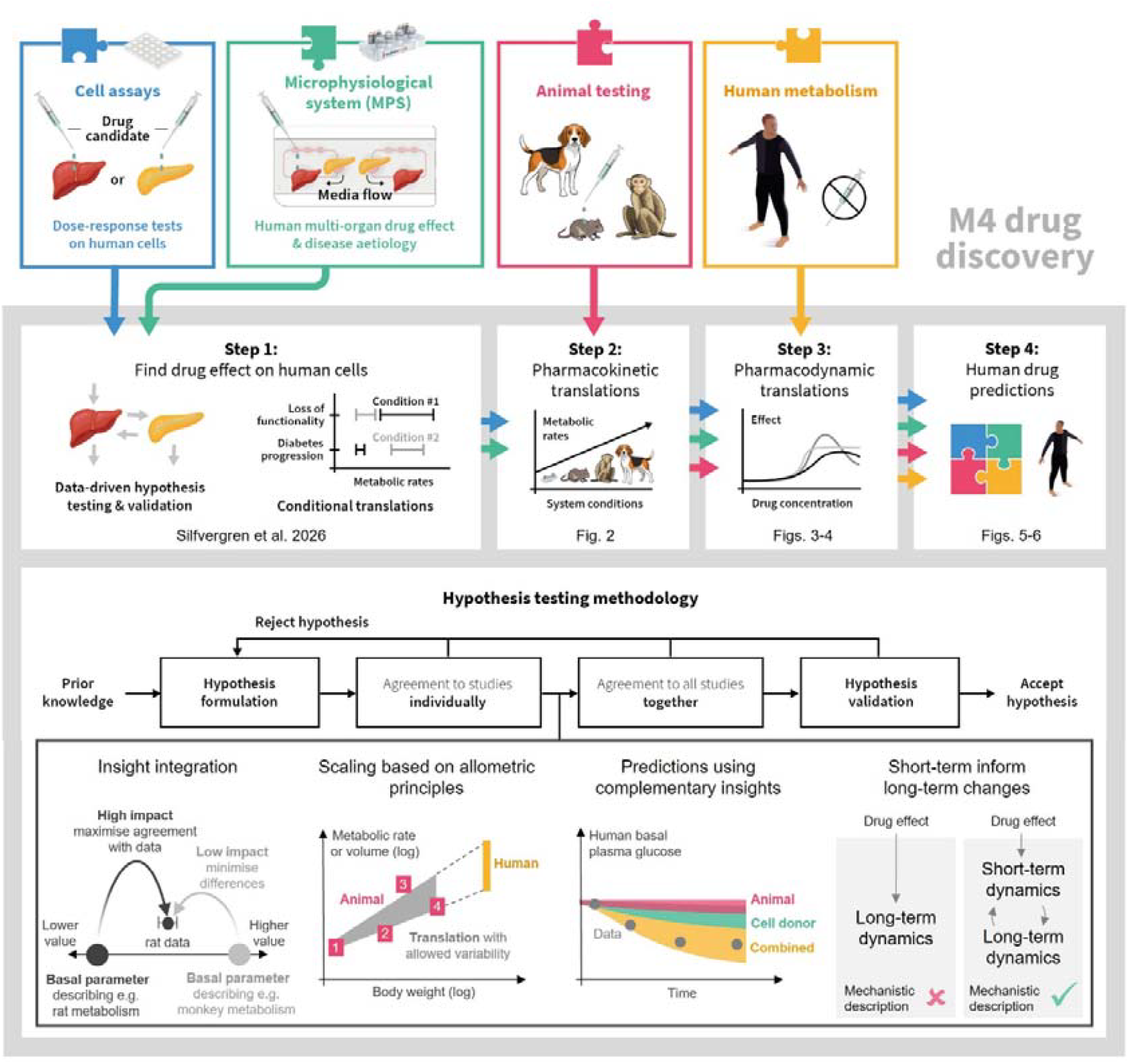
M4 drug discovery methodology. This illustration shows how M4 drug discovery integrates information from cell assays (blue), microphysiological systems (MPS) (turquoise), animal studies (pink), and prior clinical human data, e.g., baseline data without drug responses (yellow) to generate human drug predictions (top grey panel). The methodology uses data-driven hypothesis testing, where a model structure represents a biological hypothesis, to develop an M4 model (a mathematical model with Mechanistic, Multi-timescale, Multi-level, and Multi-species properties). The model-based hypothesis is tested for its ability to jointly describe diverse studies and is validated by predicting new, independent data.

## RESULTS

We illustrate M4 drug discovery by retrospectively performing it on the GLP-1RA exenatide (**Fig. 1**). An M4 model specifies a minimal set of biological and pharmacological mechanisms that jointly explain observations across many studies, integrating all available evidence into a single coherent analysis rather than treating each study in isolation. Integration is achieved through four elements in the hypothesis testing workflow (**Fig. 1**, bottom grey panel). First, **insight integration**, where studies are modelled together by sharing parameters that reflect common biology, while constraining the variability of the parameters to only variations that are needed to capture differences; this enables information integration across all conditions and studies. Second, **allometric scaling**, in which systematic differences between studies (for example, species or body-size effects) are represented with scaling functions (grey region) and complemented by permitted deviations (boxes) for study-specific effects not explained by scaling. Third, **complementary insights**, where observations across preclinical systems (e.g., multi-species pharmacokinetics, drug dose-responses from human cells, and additional effects observed in animal studies), are combined to produce clinical predictions. Fourth, **short-term informed long-term changes**, where the mechanistic underpinning of drug effect is physiologically described by modelling both short- and long-term changes, in which short-term dynamics propagate into cumulative long-term changes.

The M4 drug discovery of exenatide is exemplified in four steps (**Fig. 1**, steps 1-4). In our previous work (11), we completed step 1 by establishing the M4 model foundation that links exenatide’s effects from cell-based assays through MPS, using 3D human donor primary pancreatic microtissues (islets) and the EndoC-βH5 pancreatic cell line. Building on that foundation, we now complete the M4 drug discovery for exenatide by first establishing a pharmacokinetic translation to humans (step 2), then the pharmacodynamic translation to humans (step 3), and finally leveraging all prior steps to predict clinical outcomes (step 4).

### Establishing pharmacokinetic translation

The second of the four steps in the M4 drug discovery exemplification translates the pharmacokinetics of exenatide to humans (**Fig. 2**). We developed a pharmacokinetic model (**Fig. 2A**) describing drug absorption, distribution, and elimination and scaled the model based on allometric principles (**Fig. 2A**, dotted lines). The final model consists of 23 parameters (10 representing pharmacokinetic parameters and scaling, and 13 describing variability). The model structure and parameter values are detailed in the supplementary material (**Note S1**).

**Figure 2.**
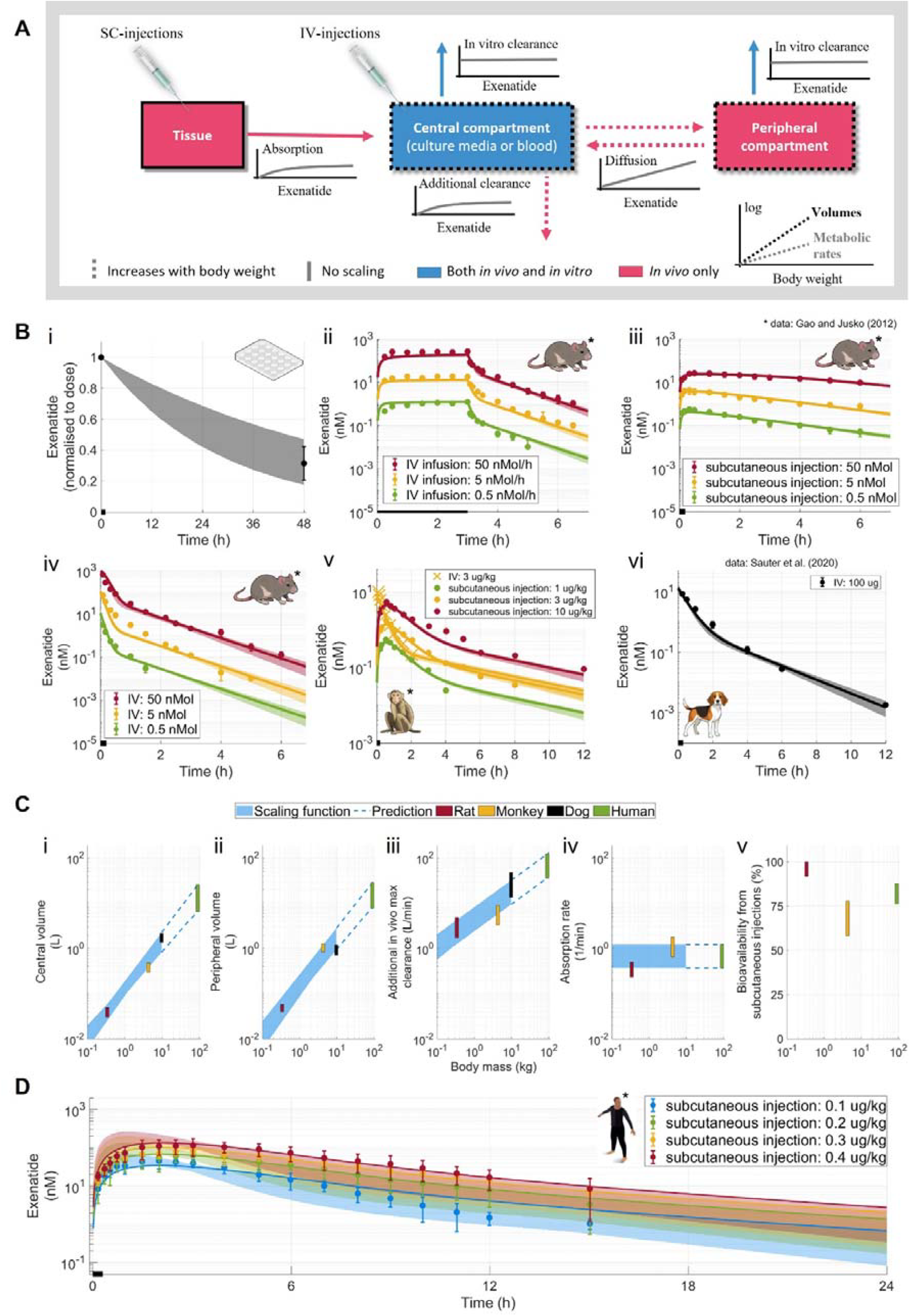
Pharmacokinetic model and agreement to exenatide exposures. **(A)** Two-compartmental pharmacokinetic model structure. **(B)** Pharmacokinetic data and model agreement to all estimation data jointly. **i)** Human cell culture studies normalised to initial dose (Table S1). **ii)** Intravenous (IV) infusion in rats (22). **iii)** Subcutaneous injections (SC) in rats (22).**iv)** IV infusions in rats (22). **v)** Exenatide exposure in monkeys (22): IV injections (crosses) and SC injections (circles). **vi)** IV injections in dogs (29). Lines represent the pharmacokinetic model’s best fit to all estimation data (Fig. 2B). **(C)** Pharmacokinetic translation of metabolic rates and volumes (y-axis) relative to body weight (x-axis). **i)** Central volume. **ii)** Peripheral volume. **iii)** Additional in vivo max clearance. **iv)** Absorption rate. **v)** Bioavailability from SC-injections. **(D)** Pharmacokinetic human data and model predictions of four SC injections (22). Lines represent the pharmacokinetic model’s best prediction from all estimation data (Fig. 2B). For all panels: coloured areas indicate model uncertainty; error-bars show population data (mean ± SD); black lines on x-axis show drug administrations.

The pharmacokinetic model was trained to describe exenatide pharmacokinetic data from our cell cultures, rats (22), non-human primates (NHP) (22), and dogs (29) (**Fig. 2B**). In cell cultures, several *in vivo* clearance mechanisms are absent (e.g., renal clearance and, to a large extent, metabolic clearance), which results in a long half-life (**Fig. 2B i**), in line with the general understanding of therapeutic peptides (23). We found that the *in vitro* clearance across all cell culture studies (T**able S1**) could be described by one-compartment kinetics (**Fig. 2A**, blue) with a half-life of approximately 30 h (uncertainty range: 19–44 h). This is similar to our studies performed without cells (**Table S2**) and previous reports of exenatide’s in vitro degradation (30,31). To represent pharmacokinetics in animals, we extended the model to two-compartment kinetics and added a second clearance pathway to represent *in vivo* processes absent *in vitro* (**Fig. 2A**, red). We then fitted the model to all preclinical pharmacokinetic datasets simultaneously, spanning seven doses, three administration routes, and four experimental systems (cells and three animal species). This simultaneous analysis yields a unified, better-informed conclusion than separate, study-by-study fits and is possible due to the <u>M</u>ulti-species properties of the pharmacokinetic model.

The pharmacokinetic translation was evaluated and compared across preclinical systems (**Fig. 2C**). We found sufficient agreement with data using a logarithmic scaling function, where volumes increased more strongly with body weight (**Figs. 2C i–ii**, slope = 0.85–1.15) than metabolic rates (**Fig. 2C iii**, slope = 0.60–0.70) (**Fig. 2C**, blue area). These observations align with previously reported values for peptides like exenatide (23,32) and the commonly assumed slopes (rates=0.75; volumes=1) sometimes referred to as Kleiber’s law (33). Deviations from the scaling functions are described using the same methodology introduced to describe variability between the cell culture studies (step 1 of M4 drug discovery), where model parameter variability is restrained but allowed to jointly describe the data (11). The resulting parameterization for each preclinical system can be viewed as the product of the allometric scaling function and a species-specific deviation term, yielding estimates for rats (**Fig. 2C**, red bar), monkeys (**Fig. 2C**, yellow bar), and dogs (**Fig. 2C**, black bar). We then used the pharmacokinetic model to generate human predictions (**Fig. 2C**, green bar). In the extrapolation, we assumed that the observed correlations of volumes and metabolic rates with body weight extend beyond the estimation range (**Fig. 2C** dotted lines). The absorption rate and bioavailability were assumed to be independent of body weight (**Figs. 2C iv–v**), which are two common assumptions in animal-to-human extrapolations (34,35). The time-dependent fate of exenatide, including its distribution across compartments and elimination through different clearance pathways, is illustrated in **Fig. S1**.

The pharmacokinetic translation was validated by successfully predicting four human SC-injection profiles (22) (**Fig. 2D)**. The agreement with the human data was supported by visual inspection and not rejected by a χ^2^-test 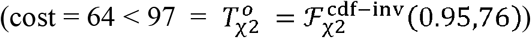.

In summary, in Step 2 we established an exenatide pharmacokinetic translation from joint insights of cell cultures and animal studies to successfully predict human SC exenatide profiles.

### Describing exenatide’s effect in rats

The third step in the M4 drug discovery exemplification is to establish a pharmacodynamic translation. To achieve this, we constructed the first M4 model of exenatide, by merging our newly developed understanding of exenatide (steps 1-2) with our pre-existing mathematical model describing human meal responses (28), and then evaluated whether the model could describe the effect of exenatide in a previously reported rat study (22) (**Fig. 3**).

**Figure 3.**
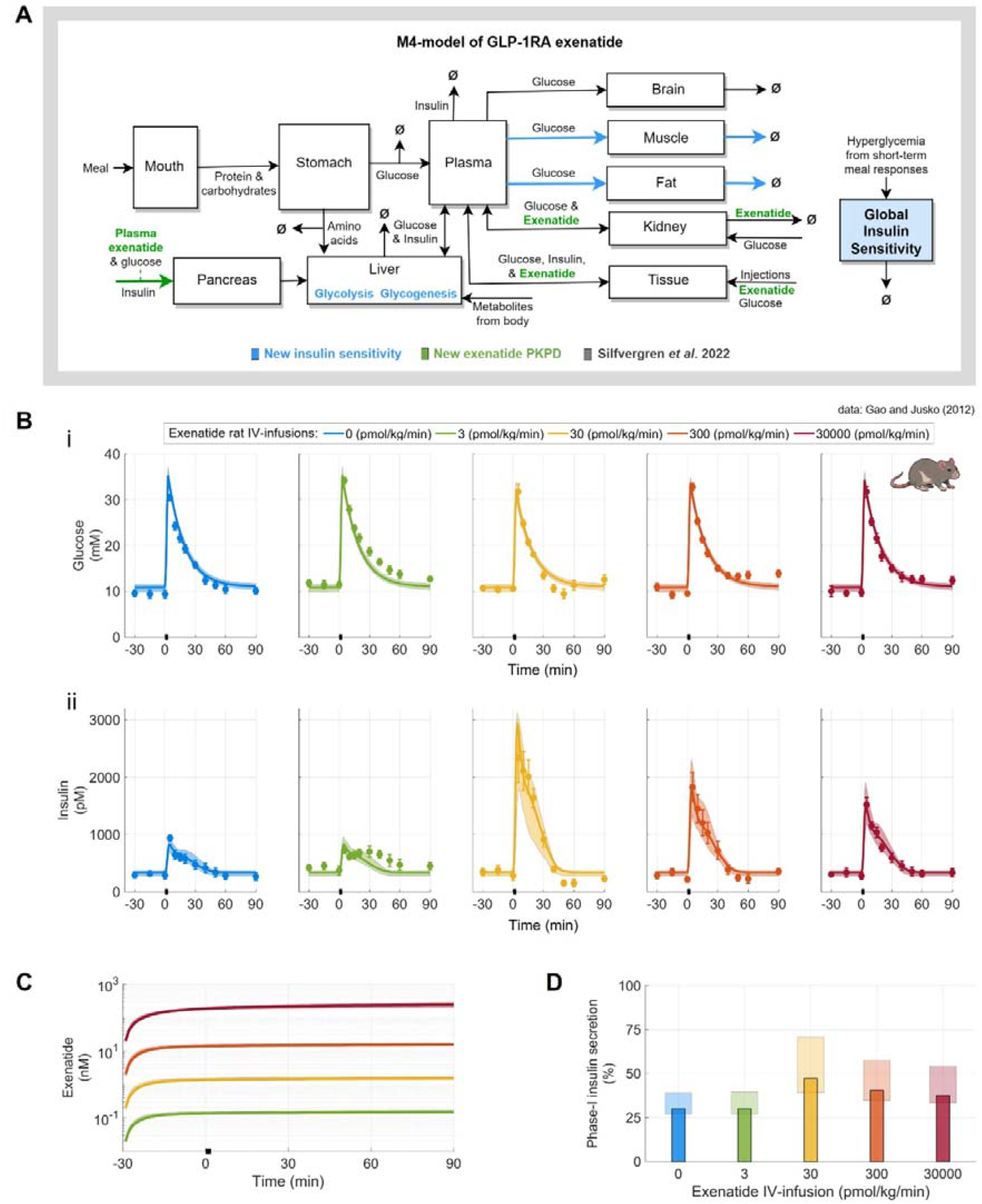
M4-model description of exenatide’s effect in rats. **(A)** Schematic of the M4 model describing whole-body metabolism affected by the GLP-1RA exenatide: new description of exenatide’s pharmacodynamics and pharmacokinetics (PKPD; green), and a new insulin sensitivity component (blue), are integrated with a pre-existing model (28) developed for clinical meal responses in the absence of drug (grey). Squares denote different parts of the body; arrows illustrate metabolic flows and rates; Ø indicates an outflow from the model. **(B)** Model agreement with data of rat glucose□challenge experiments (22). **i)** Plasma glucose concentrations. **ii)** Plasma insulin concentrations. (**C)** Predicted exenatide concentrations of the four IV infusions. (**D)** Predicted contribution of phase-I insulin secretion to total insulin secretion. Phase-I insulin secretion was defined as the first 10 minutes after glucose injection (**Note S1**). For all panels: Lines and bars represent the M4-model’s best description of all estimation data (pharmacokinetics, Fig. 2B; and pharmacodynamics, Fig. 3B); coloured areas indicate M4-model uncertainty; error bars show population data (mean ± SD); black bars along x-axis indicate timepoint for glucose injection.

The M4 model of exenatide was constructed to integrate pharmacology with whole□body glucose regulation (**Fig. 3A**). The model extends our pre-existing mathematical model of human meal responses (28) by incorporating exenatide pharmacodynamics (step 1) and pharmacokinetics (step 2) (**Fig. 3A**, green), and by adding a dynamic insulin sensitivity component (**Fig. 3A**, blue). Within the model, exenatide enhances glucose-dependent insulin secretion, thereby amplifying insulin-dependent processes including glucose uptake in muscle and adipose tissue, hepatic glycolysis and hepatic glucogenesis. Importantly, these insulin-dependent processes are governed by a global insulin sensitivity parameter that slowly increases when postprandial plasma glucose concentration is elevated and gradually decreases over time, providing a mechanism by which short □term meal and drug effects can drive longer □term metabolic adaptations. The full model consists of 72 parameters describing a simplified representation of whole-body glucose metabolism and exenatide pharmacology. The detailed model structure and parameter values are provided in the supplementary material (**Note S1**). The original human meal response model was designed and trained to describe clinical data in the absence of drug treatment. To adapt it from human meal responses to rat glucose injection-responses, we employed a minimal adaptation strategy: a) all parameters were refitted to the rat dataset, and b) the insulin production driven by the rate of change of glucose was constrained to contribute only positively to the total insulin production (**Note S1**).

The M4 model reproduced the observed effect of exenatide on rat plasma insulin secretion following 5.7 mmol/kg glucose injections 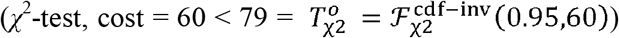 (**Fig. 3B**). In the underlying study, five exenatide IV infusions were initiated 30 min before the glucose injections, but exenatide exposure was not reported. We therefore used the previously established pharmacokinetic model (step 2) to predict exposures (**Fig. 3C**), yielding concentrations consistent with the original study (22). The maximal effect of exenatide on insulin secretion (**Fig. 3B ii**) was predicted at approximately 1.5 nM (**Fig. 3C**, yellow). This estimation was enabled by the model’s <u>M</u>ulti-species and <u>M</u>echanistic structure, which links pharmacokinetics, pharmacodynamics, and systems glucose regulation.

The proportion of phase-1 insulin secretion was predicted at approximately 30% in the non-treated group (**Fig. 3D** blue, uncertainty range: 26–40 %) and increased with exenatide treatment (**Fig. 3D**; blue compared to non-blue). Estimating exenatide’s effect on biphasic insulin secretion was made possible by the M4-model’s <u>M</u>ulti-level structure.

### Translating the pharmacodynamics of exenatide to humans

The third part of M4 drug discovery was completed by establishing a pharmacodynamic translation strategy from preclinical systems to humans (**Fig. 4**). This strategy integrated exenatide dose–response data from rats and human cell cultures, a potency calibration step, and an appetite effect identified in a prior rat study.

**Figure 4.**
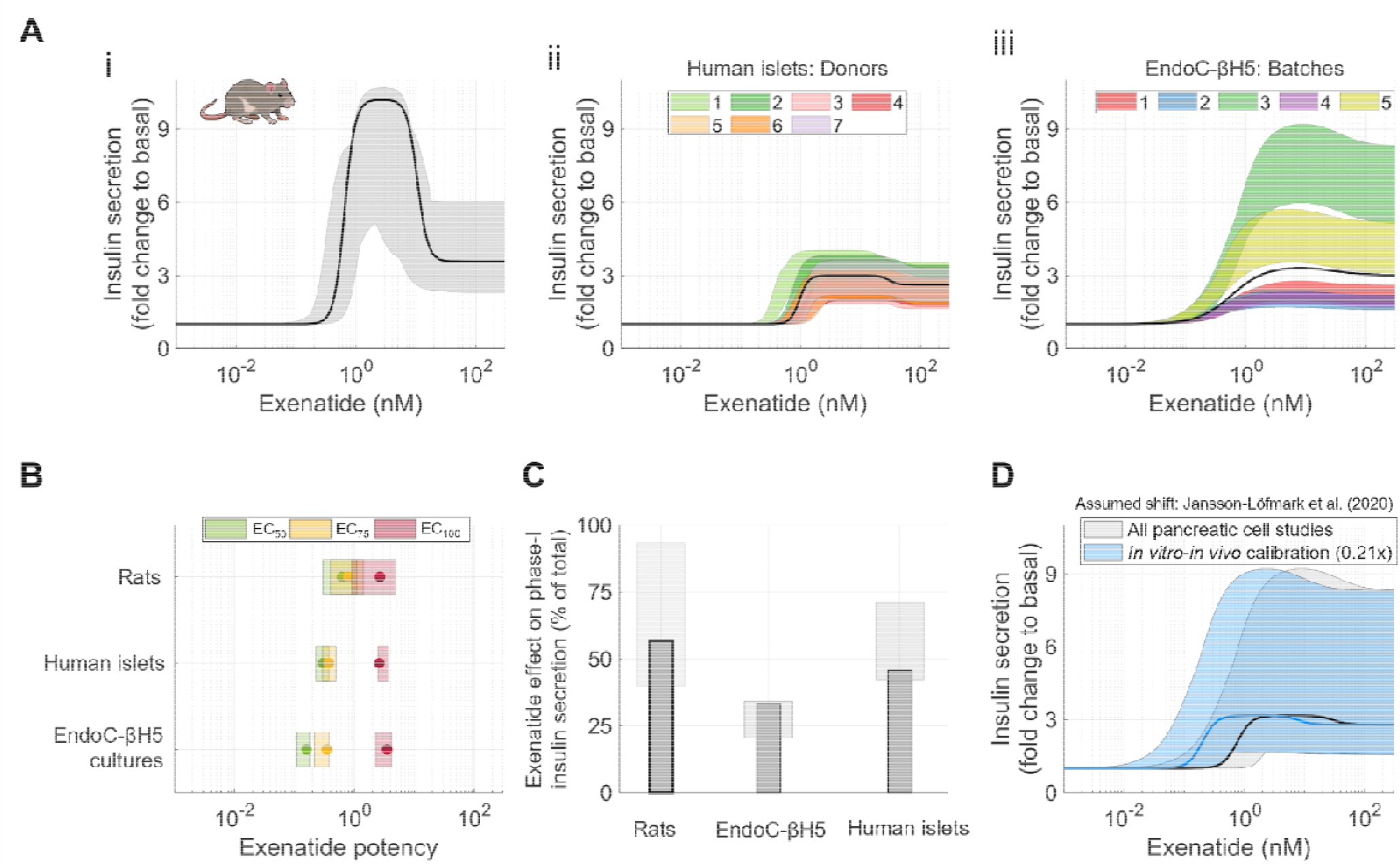
Components of the pharmacodynamic translation strategy for exenatide. **(A)** Predicted exenatide dose-response curves. **i)** Rats. **ii)** Human islets, with human donor-specific responses by colour. **iii)** EndoC-βH5 batches, with batch-specific responses by colour. **(B)** Exenatide potency comparisons between rats, human islets, and EndoC-βH5. Dots represent best agreement to data. **(C)** Predicted fraction of exenatide effect on phase-I insulin secretion relative to the total exenatide effect on insulin secretion. **(D)** Human pancreatic dose-response curves without calibration (black) and after calibration (blue), based on preexisting historical in vitro–in vivo shift data (36). For all panels: Lines and bars represent the M4-model’s best description of all estimation data; coloured areas indicate model uncertainty.

First, we compared the effect of exenatide on insulin secretion across studies in terms of dose-response shapes (**Fig. 4A)**, potency (**Fig. 4B**), and contributions to its biphasic behaviour (**Fig. 4C**). We made four key observations: a) a bell-shaped exenatide dose-response curve was evident across all systems (**Fig. 4A**); b) the cell-culture-derived effect of exenatide on glucose-dependent insulin secretion (11) was also apparent in the rat study (**Fig. 3A ii**; -30 to 0 min); c) the best-fit of rat EC_50_ and EC_75_ exceeded those from both pancreatic cell models (**Fig. 4B**, green and yellow dot); and d) exenatide influenced both phases of insulin secretion in all preclinical systems evaluated (**Fig. 4C**).

Second, we calibrated potency to human *in vitro* measurements of exenatide using published *in vitro*– *in vivo* potency shift data from a broad analysis of small-molecule drugs (36). That study quantified potency shifts across predefined drug categories. Although exenatide is a peptide rather than a small molecule, we selected the categories judged most relevant to its pharmacology and disposition as the closest available approximation. Specifically, we used the mean shift across the categories alimentary tract and metabolism (0.15x), G protein-coupled receptor agonists (0.23x), extracellular (0.25x) and no reported active metabolites (0.22x), yielding an estimated 0.21x reduction in EC_100_ and the adjusted dose–response curve shown in Fig. 4D.

Finally, we incorporated an effect of exenatide on appetite into the pharmacodynamic translation strategy based on a prior rat study (37). Using a minimal approach, exenatide was assumed to reduce daily caloric intake by a fixed 30% once its effect on insulin secretion reached 10% of maximal efficacy (**Note S1**).

In summary, Step 3 established an M4 model that describes the effect of exenatide in rats, and defined a human pharmacodynamic translation strategy informed by cell-culture and rat data.

### Evaluating simulation alternatives to predict human exenatide treatment outcomes

In the final part of the M4 drug discovery exemplification, we integrated all prior steps (Steps 1–3) to perform a preclinical-to-clinical extrapolation of exenatide. Specifically, we simulated a 30-week human study of a long-acting exenatide formulation administered once weekly at 2 mg (38) (**Fig. 5**). The predicted outcomes focused on long-term changes in basal plasma glucose, inferred from short-term meal-induced and exenatide-induced responses. The simulated diet provided 2700 kcal/day (55% carbohydrates, 15% protein, and 30% fat), divided equally among breakfast (starting at 07:00), lunch (13:00), and dinner (18:00). All meals were assumed to be consumed at a constant rate over 15 minutes. Further details of the simulated diet are included in the supplementary material (**Note S1**).

**Figure 5.**
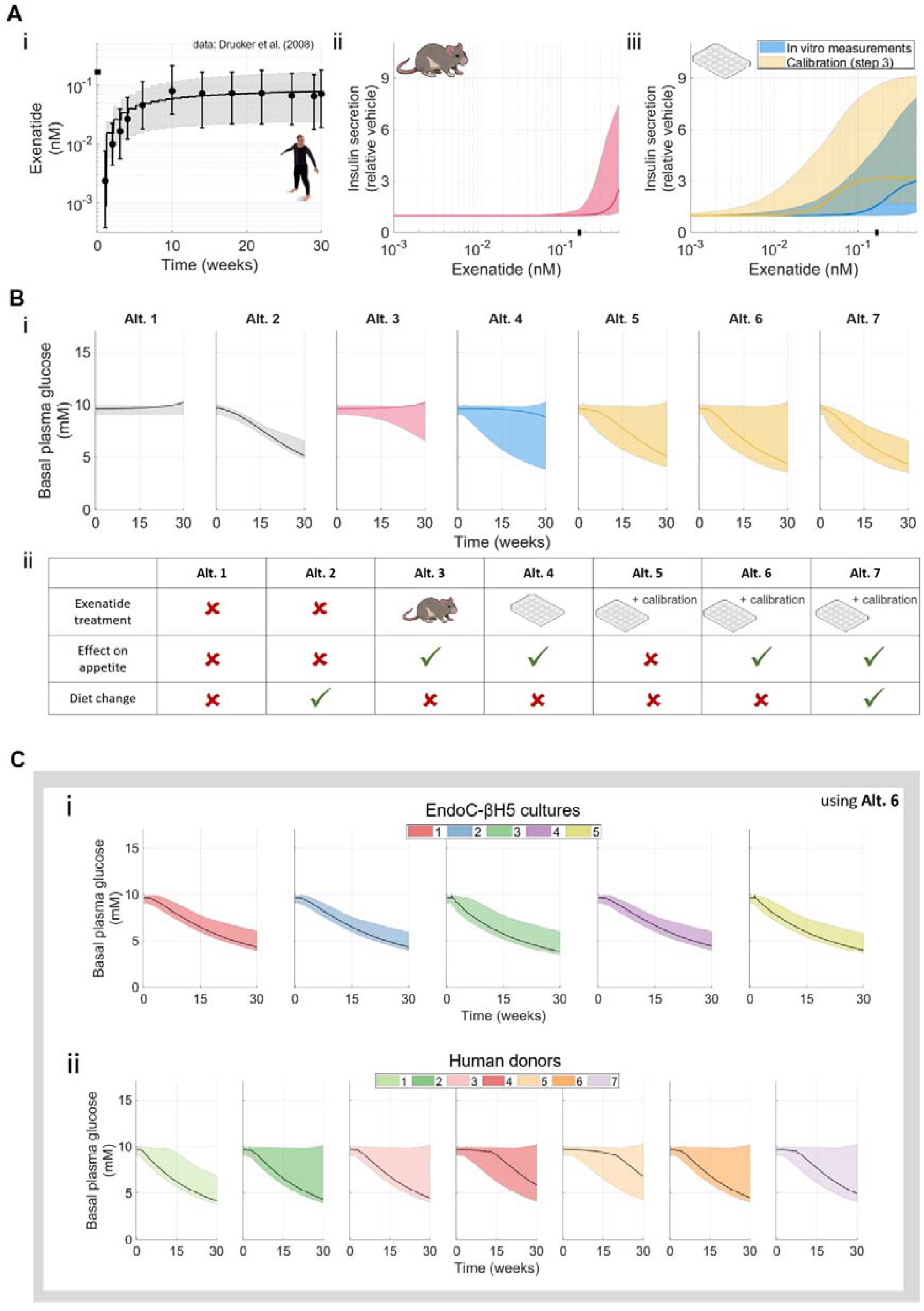
M4-model simulation of a 30-week exenatide treatment with different combinations of preclinical insights. **(A)** M4-model description of exposure and dose-response relationships. Black bar represents maximum simulated exenatide exposure. **i)** Agreement to reported exposure data (38). Error bars depict mean ± interquartile (Q1–Q3) range, matching the source data. The absorption rate was calibrated for the long-acting release formulation of exenatide (Note S1). **ii)** Exenatide dose-response in rats. **iii)** Exenatide dose-response of all in vitro measurements: without the potency calibration (blue) and with the potency calibration from step 3 (yellow). **(B)** Seven simulation alternatives of the 30-week human exenatide treatment study. **i)** Predicted basal plasma glucose trajectories. **ii)** Settings for each simulation alternative. **(C)** Human donor and EndoC-βH5 batch-specific predictions for the 30-week treatment using Alternative 6. **i)** Five EndoC-βH5 batches. **ii)** Seven human islet donors. For all panels: Lines represent the model’s best description of all estimation data; coloured areas indicate model uncertainty.

During the 30-week human study, plasma exenatide concentration approached steady state of approximately 0.02–0.2 nM by week 10 (**Fig. 5A i**). At this exposure, the rat □derived dose–response predicts no exenatide □induced increase in insulin secretion (**Fig. 5A b**, line), whereas the dose– response relationships derived from human cells do (**Fig. 5A c**, lines).

The 30-week human exenatide study was simulated using seven alternatives to assess how preclinical insights combine to influence outcomes (**Fig. 5B**). Three factors were varied: i) exenatide treatment on versus off, and if on, potency sourced from rats or human cell culture, with or without calibration; ii) inclusion versus exclusion of an exenatide effect on appetite; and iii) the presence versus absence of a diet change. From these simulations several observations were made. Basal plasma glucose remains stable in the absence of both diet change and exenatide treatment (**Fig. 5B**, Alt. 1). A change in diet alone can shift basal plasma glucose toward healthy levels (**Fig. 5B**, Alt. 2). When exenatide treatment is simulated using rat-derived potency without an active diet change, prediction uncertainty increases toward a healthy range, although the best description to the data still indicates no change (**Fig. 5B**, Alt. 3 compared with Alt. 1). Replacing rat-derived potency with human cell-culture-derived potency further broadens the uncertainty toward healthy levels, and the best description to the data shows a small improvement in basal plasma glucose (**Fig. 5B**, Alt. 4). When alternative 4 is simulated using the *in vitro*–*in vivo* potency calibration, the best agreement with data shifts substantially toward healthy levels, while model uncertainty remains largely unchanged (**Fig. 5B**, Alt. 4 compared with Alt. 6). Removing exenatide’s effect on appetite from alternative 6 reduces the predicted improvement in basal plasma glucose (**Fig. 5B**, Alt. 6 compared with Alt. 5). Finally, combining alternative 6 with a diet change markedly decreases model uncertainty towards healthy levels (**Fig. 5B**, Alt. 6 compared with Alt. 7), and the improvement in basal plasma glucose exceeds that obtained by dieting alone (**Fig. 5B**, Alt. 7 compared with Alt. 2).

Human cell donor- and cell-line batch-specific predictions of the 30-week human exenatide study were generated using simulation alternative 6 (**Fig. 5C**) and dose-response curves established in step 3 (**Fig. 4A**). We found that all EndoC-βH5 batches produced prediction intervals consistent with an improvement in basal plasma glucose (**Figs. 5C i**). The greatest glucose improvement was observed for batch 3, which also exhibited the highest exenatide efficacy (**Figs. 4A iii**, green). In comparison, human donor-to-donor variability was predicted to exceed cell-line batch-to-batch variability (**Figs. 5C ii**). For human donor 1, all simulations indicated an improvement in basal plasma glucose (**Figs. 5C ii**, area), aligning with this donor’s higher measured exenatide potency (**Figs. 4A ii**, donor 1 has the highest). Similarly, the smallest predicted improvement was from human donor 5, with a benefit first predicted once long-acting exenatide exposure reached approximately 0.05 nM (**Figs. 5C ii**, line; **Fig. 5A i**, week 10).

### Predicting a 30-week exenatide treatment with quantitative agreement to data and qualitative agreement to general aspects of human metabolism

In the final part of the M4 drug discovery exemplification, we compared the human predictions with glucose data from the 30-week human exenatide treatment study, and evaluated qualitative consistency with established features of human metabolism (**Fig. 6**).

**Figure 6.**
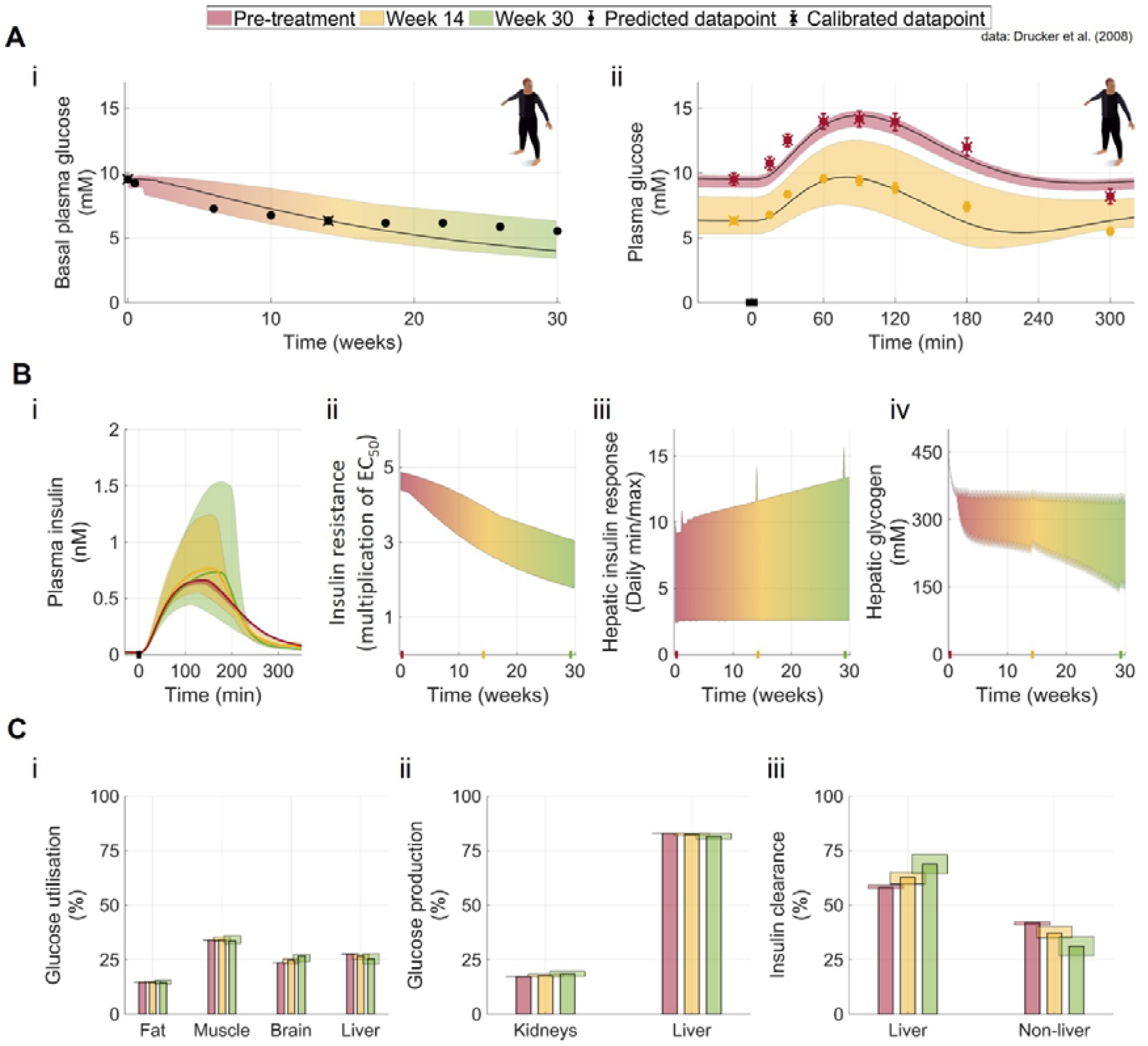
M4-model prediction for a 30-week exenatide treatment in humans. **(A)** Predicted plasma glucose concentrations compared clinical observations (38) using simulation alternative 7 (**Fig. 5B**). **i)** Basal plasma glucose concentration. **ii)** Meal-stimulated glucose responses to identical mixed-meals. **(B)** Predicted variables not measured in the 30-week study. **i)** Plasma insulin responses to the identical mixed-meals. **ii)** Insulin resistance. **iii)** Daily maximum and minimum hepatic insulin response. **iv)** Hepatic glycogen concentrations. **(C)** Predicted organ- and tissue-specific contributions to whole-body glucose metabolism. **i)** Relative glucose utilisation by fat, muscle, brain, and liver. **ii)** Relative glucose production by liver and kidney. **iii)** Relative insulin clearance by hepatic versus extrahepatic pathways. For all panels: Lines represent the model’s best description of all estimation data; coloured areas indicate model uncertainty; colours indicate treatment length (0 weeks, red; 14 weeks, yellow; 30 weeks, green).

Among the tested configurations, simulation alternative 7 provided the best overall match to the clinical data (**Fig. 6A**). This prediction was statistically acceptable, since it was not rejected by a χ^2^-test 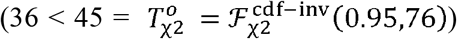, and since it passed subsequent qualitative checks against general metabolic behavior. The model predicted a marked improvement in basal plasma glucose (**Figs. 6A i**), decreasing from 9.5 mM (uncertainty range: 8.8-9.9 mM) to 4.0 mM (uncertainty range: 3.4-6.3 mM), with saturation toward normoglycemia (**Fig. 6A i**). It further reproduced the observed reduction in postprandial glucose amplitude for identical mixed meals consumed pre-treatment and after 14 weeks of treatment (**Fig. 6A ii**). Because the human study did not report mixed-meal meal size, we calibrated the M4 model to the pre-treatment response, which yielded an estimated meal of 900 kcal (**Fig. 6A ii**, red; marked with ‘x’), a physiologically plausible value (39).

To further evaluate the 30-week prediction, we simulated variables not measured in the study and compared them qualitatively to literature observations on plasma insulin, insulin resistance, hepatic insulin response, and hepatic glycogen (**Fig. 6B**).

First, the model predicted that meal □stimulated insulin responses would change over the course of exenatide treatment (**Fig. 6B i**). The best description of the study indicated a slight increase in peak postprandial insulin from approximately 650 to 750 pM (uncertainty range: 650±25 to 1175±425 pM). These predictions are in qualitatively consistent with prior reports, where increased insulin concentrations during meals are attributed to the exenatide-induced stimulation of insulin secretion (40,41), while reductions are explained by improvements in basal plasma glucose and increased insulin sensitivity (42,43). Moreover, basal insulin concentrations were predicted to remain unchanged throughout the study (**Fig. 6B i**; <0 min), in line with clinical observations (40,44,45).

Second, the model predicted a reduction in insulin resistance over the study period (**Fig. 6B ii**). The initial insulin-response EC_50_ was fitted to reproduce the basal plasma glucose preceding the first reported meal in the human study (**Fig. 6A ii**, red; marked with ‘x’). We found that a 4.3-to 4.8-fold increase in EC_50_ relative to the healthy population used to train the original meal-response model adequately described the prediabetic population (**Fig. 6B ii**, 0 weeks), a magnitude that is physiologically plausible in light of clinical observations (46–49). Over the course of treatment, EC_50_ declined to 1.7– to 3.0-fold above the healthy value, a reduction that is also physiologically plausible (50,51). The rate of decline in insulin resistance under euglycemic conditions (i.e., in the absence of hyperglycemia) was fitted to match basal plasma glucose level of the second reported meal in the human study (**Fig. 6A ii**, yellow).

Third, the model predicted an increased hepatic insulin response to identical meals, rising from approximately 10-fold basal values at baseline (**Fig. 6B iii**, 0 weeks) to approximately 18-fold by week 30 (**Fig. 6C iii**). This pattern is in qualitative agreement with human metabolism, as it arises from an elevated exenatide-induced insulin secretion (**Fig. 6B i**) and decreased insulin resistance (**Fig. 6B ii**).

Fourth, hepatic glycogen levels were predicted throughout the study (**Fig. 6B iv**). The model indicated an initial decrease during weeks 0–3, consistent with the transition to a lower-calorie diet (80% kcal/day of initial diet; described in **Note S1**). Beyond week 3, model uncertainty increases, with hepatic glycogen predicted either to stabilise at the lower level, or to decrease further towards a fasted state. Both trajectories are mechanistically plausible given the insulin- and glucose-dependency of glycogenesis. Stable glycogen levels are explained by a greater contribution from insulin-dependent reaction rates, whereas diminishing stores reflect lower insulin-dependent activity. By the end of the study, hepatic glycogen stores were predicted to lie between 150 and 350 mM, a range that is physiologically plausible in healthy and diabetic populations (52,53).

Finally, we compared organ-specific contributions to metabolic rates over 24-hour periods at three time-points: pre-treatment (red), week 14 (yellow), and end of treatment (green) (**Fig. 6C**). The predicted organ- and tissue-specific contributions to total glucose utilization **(Fig. 6C i**) were qualitatively consistent with prior studies (54). Brain glucose utilization was predicted to increase over time, reflecting its implementation as a constant-rate process, which aligns with prior modelling approaches (55). We also compared renal and hepatic glucose production, with the majority of glucose production predicted to originate from the liver (**Fig. 6C ii**), in agreement with clinical observations (52,56). Hepatic and non-hepatic contributions to total insulin clearance were evaluated (**Fig. 6C iii**), with most insulin clearance occurring in the liver, consistent with existing literature (57).

In summary of step 4, complementary preclinical insights (from cell-based assays, MPS, and animals) were combined with untreated human data (used to develop the pre-existing human meal-response model) to enable accurate multi-temporal drug predictions in humans within a single M4 model.

## DISCUSSION

Here, we present M4 drug discovery, a methodology for drug discovery (**Fig. 1**). The utility of this methodology is demonstrated using the GLP-1RA exenatide as an example. In this exemplification, data from all major preclinical systems (cell-based assays, more complex MPS, and animals) were integrated into a M4 model. A translational pharmacokinetic model of exenatide based on cell cultures, rats, NHPs, and dogs was established (**Fig. 2**), and used to successfully predict human exposure profiles (**Fig. 2D**). The model was then developed by incorporating preclinical insights for exenatide with a pre-existing model of human meal responses in the absence of drug treatment (**Fig. 3A**). The model could successfully capture the effect of exenatide in rats (**Fig. 3B**), and, together with cell culture data, a pharmacodynamic translation strategy was determined (**Fig. 4**). Together, all components of the M4 model of exenatide were combined to evaluate how complementary insights can be leveraged to predict a human 30-week exenatide treatment (**Figs. 5-6**). The human treatment predictions were supplemented by human donor-specific and cell-line batch-specific predictions, indicating the potential of this methodology for precision medicine applications (**Fig. 5C**). Among seven simulation alternatives, the best description of the human treatment was obtained by combining all generated insights (**Fig. 6A**). The simulation approach successfully described both the incremental long-term effect on basal plasma glucose (**Fig. 6A i**) and the altered short-term meal response at different time-points during the study (**Fig. 6A ii**), with qualitative agreement with general aspects of human metabolism (**Figs. 6B-C**). In conclusion, our results demonstrate that it is possible to integrate diverse data sources along the M4-axes (<u>M</u>echanistic, <u>M</u>ulti-level, <u>M</u>ulti-timescale, and <u>M</u>ulti-species) to successfully generate clinical predictions of a drug with multi-temporal mechanisms of actions.

The human prediction was possible because of the newly developed M4 model. All M4 aspects are essential in this example. The model must be **<u>M</u>**echanistic since, for example, the effect of exenatide on insulin secretion requires a description of glucose-dependent insulin secretion. The model must also be **<u>M</u>**ulti-level, since for example the two phases of insulin secretion are affected by intracellular insulin storage of readily releasable pools. Similarly, the model must be **<u>M</u>**ulti-timescale, since the effect of exenatide on meal responses occurs over minutes which, over time, propagates long-term effects such as prevention of type 2 diabetes. Finally, the model must have **<u>M</u>**ulti-species properties, because complementary insights arise from animal experiments and human cell studies. Similarly, all preclinical insights generated across all parts of M4 drug discovery are essential in this example. For instance, both an appropriate pharmacokinetic response (**Fig. 2**) and exenatide’s effect on appetite were only observable in animals (37). In contrast, the most accurate human exenatide dose-responses were obtained from human cell studies (**Fig. 5**). Notably, the **<u>M</u>**ulti-species properties of this example go beyond strict animal species differences, where data are also incorporated from different human donors and cell-line batches (**Fig. 4A**). With an M4 model, we were able to incorporate data from all these sources (Step 1 (11); **Figs. 2-3**, step 2; **Figs. 4-5**, step 3), account for and understand mechanistic differences (**Figs. 2–5**), and integrate insights into a coherent systems-level understanding (**Figs. 6–7**).

Mathematical modelling is a routine component of drug discovery, and our work builds on this tradition with M4 drug discovery. For the exenatide example, we use allometric scaling (23,58) and compartmental models (22,59) on the pharmacokinetic side. This framework can naturally be extended to physiologically based pharmacokinetic (PBPK) models (60–62). Related efforts integrate short-term (minutes) and long-term (months) dynamics, as well as cross-species pharmacokinetics and pharmacodynamics (25), aligning with the M4 concept. What distinguishes our implementation is the explicit integration of data from cell-based assays, MPS, and animal systems. This design enables accurate propagation of short-term dynamics into cumulative long-term pharmacodynamic changes, providing the mechanistic basis needed for drugs with multi-temporal mechanisms of action that include multiple feedback loops. In this way, the M4 approach mitigates persistent hurdles in preclinical-to-clinical extrapolation for drugs such as exenatide (3–6), and provides a path to more reliable translation.

The M4 drug discovery methodology relies on a central assumption, and the overall demonstration herein could be further improved in two key ways. The central assumption lies in the choice of mechanisms and equations, where it is important to note that we do not prove that our mechanisms and assumptions are the correct ones. A first key improvement would be to refine the *in vitro*-*in vivo* potency calibration. Although the human *in vitro-in vivo* potency shifts are documented across many molecules (36), future work should aim to mechanistically explain these shifts based on receptor abundance, microenvironment, receptor turnover, and related factors (19,63). The applied calibration was derived from small-molecule drugs, and its relevance for exenatide (with a molecular weight of 4187 g/mol) remains uncertain. Three alternative potency calibration hypotheses were evaluated: calibrate rat potency readings using historical rat *in vitro-in vivo* potency shifts; calibrate human cell potency readings with the same rat *in vitro-in vivo* potency shifts; and calibrate rat potency using historical rat-to-human potency shifts. Nonetheless, all three alternatives were deemed less plausible than the tested approach due to large compound-specific variability (64–66). The second key improvement would be that future M4 implementations should aim to use all information generated during preclinical drug development for clinical predictions. Herein, exenatide pharmacokinetics, effects on insulin secretion and appetite, dose–response relationships, and hepatic insulin response were integrated into a human metabolism model developed in the absence of drug treatments (**Fig. 3A**). We argue that future M4 development should strive toward a complete translation, using either both human and preclinical data in model training, or a full extrapolation without the reliance on human data.

In conclusion, we present a drug discovery methodology that enables human drug predictions from fundamentally different preclinical systems. We demonstrate how a single quantitative framework can generate person-specific clinical predictions from cell donor data and how complementary insights from different systems can strengthen clinical predictions. The approach enables preclinical-to-clinical extrapolations especially suited for compounds with multi-timescale mechanisms of actions, strengthening preclinical drug discovery.

## METHODS

The general mathematical modelling follows a data-driven hypothesis-testing approach which has been described in previous works (67,68). The general methodology is, in brief, described below. A detailed description of the M4 model and how it was used in all parts of our analysis is included in the supplementary material (**Note S1**).

The mathematical model is constructed using ordinary differential equations (ODEs) and follows the same general formulation (**Eqs. 1A-C**).

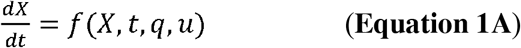

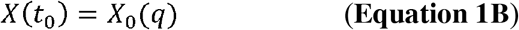

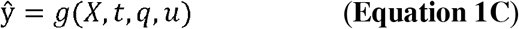

where *f* and *g* are smooth non-linear functions dependent on: *X* which is a vector of state variables, for example, concentrations of metabolites in specific compartments; *q* which is a vector of model parameters, representing for example metabolic rates and scaling constants; *u* which is a vector of input signals, for example meals and drug administration; *X*(*t*_0_) which is a vector of initial conditions *X*_0_(*q*)that are declared from model parameters *q*; ŷ which is a vector of model outputs, for example metabolites in media, insulin sensitivity, etc.

### Parameter estimation

The agreement between the model simulation and experimental data was evaluated using sum-weighted least squares (**Eq. 2**):

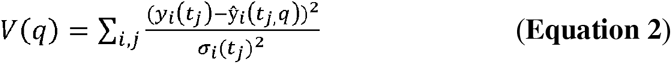

where _*y*_*i*_ (*t*_*j*_) is the measured mean of a group of datapoints; where *j* is a specific datapoint within the experimental setup *i* at the time-point *t*; where σ _*i*_ (*t*_*j*_)is the standard deviation of the group of datapoints; where ŷ_*i*_(*t*_*j*,_*q*) is the mathematical model equation value. To minimise model overfitting to small standard deviations, the mean σ_*i*_ (*t*_*j*_)_ of the group was used. The weighted sum of residuals *V*(*q*) was minimised in an optimisation problem to find simulations with the best agreement to data.

MATLAB R2019b (Natick, Massachusetts: The MathWorks Inc) was used for all parts of the mathematical modelling including simulations, plotting, and parameter estimations. The mathematical model was implemented in MATLAB using IQM toolbox (69). For the parameter estimation, we employed two distinct optimization algorithms: a) particle swarm (MATLAB global optimisation toolbox); b) PESTO Monte Carlo sampling (70). The parameter estimation was done with 16 parallel workers (MATLAB Parallel Computing Toolbox) with a stopping criterion defined by a maximum number of stall iterations (700) and a function tolerance (10^−5^). Parameter uncertainty was assessed from a collection of statistically non-rejected parameter sets during global and local refinement.

A statistical χ^2^-test was performed to quantitatively evaluate a model’s agreement to data. The statistical test was done with the null hypothesis that the experimental data had been generated by the model, assuming that the experimental noise was additive, independent, and normally distributed (68). In practice, the agreement to data was rejected if the weighted residuals *V*(*q*) had a higher value than the χ ^2^ -threshold. This is given by the inverse χ ^2^cumulative density function (**Eq. 3**):

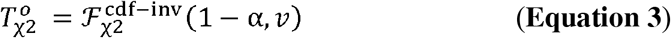

Where 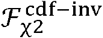 is the inverse density function; α is the significance level (α = 0.05 was used); where *v* is the degrees of freedom, which was equal to the number of data points in the dataset. In addition to statistical χ^2^-tests, model rejection was made through qualitative assessments. These qualitative assessments were implemented into the objective function where the output describes the agreement to both qualitative dynamics and data (**Eq. 4**):

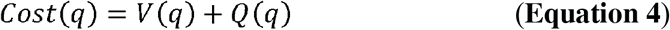

where *Cost* is the objective function that is minimised in the parameter estimation with parameters q; where*V*(*q*) is model agreement to data; where *Q*(*q*) is a function returning a scalar value based on the model agreement to qualitative system dynamics. Residuals of subject-specific data were taken into consideration in the qualitative assessment term due to a lack of standard deviation *σ*_*i*_ (*t*_*j*_)

### Allowing constrained parameter variability

The pharmacokinetic translation across animals was developed by allowing constrained variability in the same methodology as previously described (11). In brief, model parameters are structured in two layers: a) a base parameter, which describes a reaction applicable to all conditions (e.g. drug clearance); and b) a condition alteration, which describes a change in the reaction from the base parameter to a specific condition (e.g. rat-specific drug clearance). The difference between the base parameter and the condition-specific parameter is referred to as a parameter adjustment step (11), which was calculated in the objective function (**Eq. 5A**):

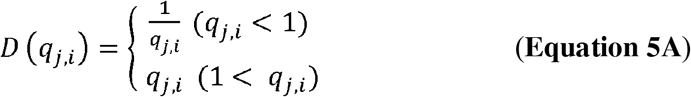

where *D*(*q*_*j,i*_) is the parameter adjustment step; where *q*_*j,i*_ is the condition alteration; where *j* indicates which base parameter is altered; where *i* specifies the condition that changes the base parameter. The adjustment step is minimised with a penalty-term in the objective function (**Eq. 5B**):

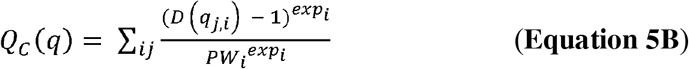

where *Q*_*c*_(*q*) is a penalty term that regularises condition-specific alterations towards 1; where 1 denotes no change to the base parameter because condition alterations act multiplicatively on the base parameter; where *PW*_*i*_ is the penalty weight for condition i; and where *exp*_*i*_ is an exponent used to scale the impact of the constraint on the objective function. The result of the minimisation is that differences between conditions are reduced, enabling the integration of insights across conditions. The parameter *PW*_*i*_ determines the strength of the penalty term: a high value implies that parameter estimation is primarily driven by agreement with the data, whereas a low value allows agreement with the data to be traded for increased insight translation between conditions. In practice, the constrained parameter variability for the pharmacokinetic translation was implemented using over various parameters repeated for each animal-type studied: a) maximum additional *in vivo* clearance (a total of 3 parameters); b) central compartment volume (a total of 3 parameters); c) peripheral volume compartment (a total of 3 parameters); d) maximum adsorption rate from subcutaneous injections (a total of two parameters; SC-injections were studied in two animals); and e) bioavailability from SC-injections (a total of two parameters). The settings used to minimise the parameter variability was *PW*_*i*_ = 0.5 and *exp*_*i*_ = 2 (**Eq. 5B**).

A detailed description of the M4 model and how it was used in all parts of our analysis is included in the supplementary material (**Note S1**).

## DATA AVAILABILITY

Data of exenatide effect in human cell culture studies were downloaded from the supplementary material of our previous exenatide-related analysis (11). The data used to describe exenatide pharmacokinetics across these studies are included in the supplementary material (**Tables S1-2**). All data of animals and humans (22,29,38) used in our analysis were collected from literature using a publicly available data extraction software (71). The digitised data is included in the supplementary material (**Tables S3-8**). In addition to the supplementary materials, all data can be downloaded from our GitHub repository (https://github.com/OscarSilfvergren/M4-drug-discovery-exenatide).

## CODE AVAILABILITY

All code used in the analysis can be downloaded from the GitHub repository (https://github.com/OscarSilfvergren/M4-drug-discovery-exenatide).

## ACKNOWLEDGEMENTS

**Johanna Dietzfelbinger** (TissUse) for figure visualisation.

The **MPS network** (AstraZeneca) for feedback and discussions over the years the analysis was done.

## FUNDING

Swedish Research Council 2023–03186 (GC)

Swedish Research Council 2023–05460 (GC)

X-HiDE, Knowledge Foundation 20200017 (GC)

CENIIT 15.09 (GC)

The Horizon Europe project STRATIF-AI 101080875 (GC)

The Swedish Fund for Research without Animal Experiments F2019-0010 (GC)

ELLIIT 2020-A12 (GC)

The County Council of Östergötland RÖ-1001928 (GC)

VINNOVA VisualSweden 2020-04711 (GC)

## AUTHOR INFORMATION

### Authors and Affiliations

**Linköping University, Linköping, Sweden**

Oscar Silfvergren, Christian Simonsson, Peter Gennemark, and Gunnar Cedersund

**SUND sound medical decisions, Linköping, Sweden**

Oscar Silfvergren and Gunnar Cedersund

**TissUse GmbH, Berlin, Germany**

Sophie Rigal, Katharina Schimek, and Uwe Marx

**German Federal Institute for Risk Assessment (BfR), Berlin, Germany**

Sophie Rigal

**Technische Universität Berlin, Berlin, Germany**

Katharina Schimek

**AstraZeneca, Gothenburg, Sweden**

Kajsa P Kanebratt, Liisa Vilén, and Peter Gennemark

**InSphero AG, Schlieren, Switzerland**

Felix Forschler and Burcak Yesildag

**Örebro University, Örebro, Sweden**

Gunnar Cedersund

### Contributions

**Conceptualization**: OS, CS, LV, PG, GC

**Methodology**: OS, CS, PG, GC

**Investigation**: OS, SR, KS, KPK, FF, BY, LV, PG, GC

**Visualization**: OS, SR, KS, LV, PG, GC

**Funding acquisition**: GC

**Project administration**: LV, PG, GC

**Supervision**: BY, UM, LV, PG, GC

**Writing – original draft**: OS, SR, KS, CS, LV, PG, GC

**Writing – review & editing**: OS, SR, KS, CS, KPK, FF, BY, UM, LV, PG, GC

### Corresponding author

Correspondence to Gunnar Cedersund.

## SUPPLEMENTARY MATERIALS

**Table S1**: Exenatide loss during cell cultures.

**Table S2**: Supplementary exenatide degradation study preformed in experimental systems without cells

**Tables S2 to S8**: Digitalised data from literature used herein.

**Figure S1:** Time-continuous pharmacokinetic differences predicted between animals and cells from identical IV infusions.

**Note S1**: Detailed description of the M4 model and how it was used in the M4 drug discovery exemplification.

**Figure S1.**
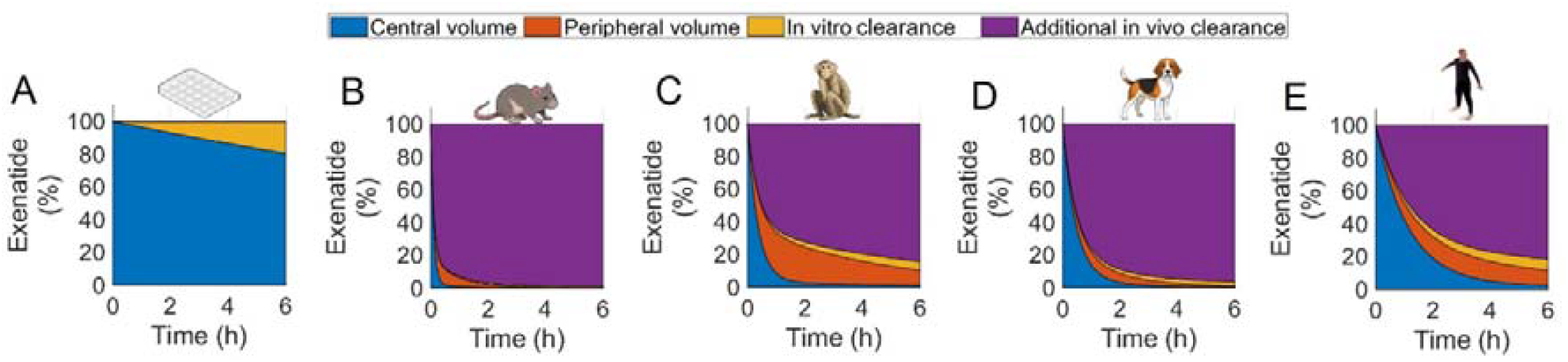
Supplementary predictions of time-continuous pharmacokinetic differences between animals and cell cultures. (A) The cell cultures (11). (B) Rats. (C) Non-human primates. (D) Dogs. (E) Humans. All panels: colours indicate exenatide distribution or loss (central volume, blue; peripheral volume, orange; in vitro clearance, yellow; additional in vivo clearance, magenta); simulations were made with best agreement to all pharmacokinetic data; all simulations were made of 1 nMol simulated either as an IV injection (animals) or directly into the media (cell cultures).

