## Supplementary material for "M4 drug discovery: human drug predictions from integrated preclinical insights exemplified with a GLP-1R agonist": Note S1

### **Supplementary Note S1**

This supplementary note provides additional information regarding the mathematical modelling. First, a step-specific description is provided of how the mathematical modelling was used in the M4 translation of exenatide. Finally, a detailed description of the M4 model is provided.

**Step-specific description of the exemplified M4 drug discovery**

###### In this section, supplementary details are provided for all steps in the M4 drug discovery exemplification of GLP-1RA exenatide.

###### Step 2: Description of pharmacokinetic translation

As described in the main text, the pharmacokinetic model was developed to describe and integrate data across preclinical systems. To integrate data across preclinical systems, metabolic rates and volumes were scaled based on body weight.

The data point used to evaluate model agreement to human cell data was calculated from all human cell studies (**Fig. 2B i**, N=115). This calculation is included in the supplementary material (**Supplementary Data S1**). In this supplementary table, the data are further analysed and divided by both measurement sequence and experimental system. We found that there were insufficient data to conclude any impact from either the measurement sequence or experimental system. An additional study was conducted to evaluate exenatide degradation without any cells in the system (**Supplementary Data S2**), which supported the assumption (main text) that exenatide in vitro clearance could be described as a constant half-life, similar to degradation. The additional study (**Supplementary Data S2**) was performed in both the same MPS system (Chip2) used to evaluate exenatide effect on liver-pancreas metabolism, and low-protein-binding tubes at 37 degrees Celsius without any spheroids/islets in the system.

In the second step of M4 drug discovery of exenatide, a comparison was preformed across preclinical systems and human (**Fig. S1**). The comparison was performed using the same dose across all systems (1 nmol IV-injections) and the declared body weights used in all pharmacokinetic simulations were in accordance with the reported values in the literature from which the data originated: rats (0.34 kg) (1), NHPs (4.31 kg) (1), dogs (9.725 kg) (2), and humans (91 kg) (1). The simulation of the exenatide concentration in our cell culture studies was performed using 605 µL total media volume, based on what was used in several of the MPS studies analysed (3).

*Step 3: Description of pharmacodynamic translation*

As described in the main text, a M4-model was developed to describe exenatide effect on a rat study and strategize a human pharmacodynamic translation. The M4-model is a further development of a pre-existing glucose meal-response model (4), with exenatide pharmacodynamic equations from our previous analysis of exenatide response in human cells, and the exenatide pharmacokinetic model from step 2.

In the M4-model, two equation alternatives were tested to describe the rat glucose injection study (Alt 1: **Supplementary Eq. S62a**; Alt 2: **Supplementary Eq. S62b**). Model versions with these alternatives are available on our GitHub repository (see main text). The first alternative is the same mathematical model used to simulate the 30-week human treatment study. Possibly because the original model (4) was not designed for glucose injections or to describe rat metabolism, its agreement with both glucose and insulin data did not pass visual inspection. This was determined even though a χ2-test did not reject the effect of exenatide on insulin levels (χ2-test, cost = 75 > 79 =  $T_{\chi2}^{o}=\mathcal{F}_{\chi2}^{cdf-inv}\left( 0.95,60 \right)$). To improve the model’s capabilities to describe the rat study, a minimal change was implemented. This change is that glucose rate of change dependent insulin production was constrained to be positive. With this minimal change, the agreement to data was improved (χ2-test, cost = 60 < 79 =  $T_{\chi2}^{o}=\mathcal{F}_{\chi2}^{cdf-inv}\left( 0.95,60 \right)$), and also passing our visual assessment. All rat simulations in the main text have this minimal change. All differences between the rat glucose injection groups with various exenatide intravenous infusions are assumed to be caused by the exenatide effect, meaning no adjustments for rat-to-rat variability were necessary to describe the data.

To ensure that the simulated rat metabolism was in a steady state in the beginning of the study, a standardised diet was simulated prior to the glucose injection. The standardised diet was simulated using the model by 21 repeated meals, each consumed over 15 minutes, with an 8-hour interval between meals (i.e. 3 meals/day). Each repeated meal was assumed to consist of 20 kcal (55% carbohydrates, 15% protein, and 30% fat), based on pre-existing literature (5,6) to ensure physiological relevance.

Exenatide effect on phase-I insulin secretion contribution to the overall insulin secretion was predicted in the main text (**Fig. 4D**). To estimate this, an assumption had to be made when phase-I insulin secretion transitions into phase-II insulin secretion, and when the meal-response transitions into a basal reaction rate. We assumed that phase-I insulin secretion is the first 10 minutes after the glucose injection, and phase-II insulin secretion takes place after the 10 minutes and ends 60 minutes after the glucose injection.

Four potential potency calibrations were hypothesised for the pharmacodynamic translation. The first hypothesised calibration was to evaluate if rat *in vitro*-*in vivo* potency shifts measured from pre-existing compounds could calibrate our estimated exenatide potency readings in rats, with the assumption that the potency shift could be similar in exenatide. Similarly, the second potential calibration hypothesised that the same rat *in vitro*-*in vivo* potency shifts could be applied to our human cell potency readings, with the assumption that rat *in vitro*-*in vivo* potency shifts could be similar as human *in vitro*-*in vivo* potency shifts. Historically, *in vitro* measurements can reflect in *vivo potency* by assuming identical receptor-specific parameters (e.g. binding affinity and rate) (7,8). But, the first and second hypothesized calibrations were not made, as the potency discrepancies are compound specific (9) and no universal rat-specific correlation could be concluded from literature. The third hypothesised calibration was to apply historical rat-to-human potency shifts to our estimated rat dose-response, with the assumption that rat-to-human potency readings might correlate over compounds. However, this was also rejected as systemic reviews of these potency shifts indicate large uncertainties and that the shifts are compound-specific (10,11). The final hypothesised calibration was to apply an *in vitro*-*in vivo* potency calibration to our exenatide translation based on historical human *in vitro*-*in vivo* potency shifts, which has historically shown that clinical potency is higher than in vitro measurements (12). As described in the main text, from these observations we generated a calibrated dose-response curve from our human cell studies by applying a 0.21x reduction in potency **(Fig. 4D**), computed as the mean of the categories exenatide falls under in the pre-existing work (12) (ATC-category, Alimentary tract and metabolism, 0.15x; GPCR agonists, 0.23x; Extracellular, 0.25x; No reported active metabolites, 0.22x).

An exenatide-induced reduction in appetite was observed in a rat study (13). In the rat study, exenatide-treated rats displayed an initial reduction of 60% (kcal/day) which returned to a steady state of 20% (kcal/day) reduction after a week with 10 μg/kg once daily (13). When increasing exenatide dosing to 10 μg/kg twice daily the steady state changed towards a 50% reduction in daily kcal. The food intake of the vehicle group reduced as well, which indicates complexity in estimating parameters such as appetite. Without sufficient appetite data—which could not be generated in our liver-pancreas cell culture studies—we decided to use a minimal approach where exenatide effects were normalised to the vehicle group and it was reduced to a static 30% reduction in appetite. In the pharmacodynamic translation strategy this reduction in appetite was incorporated with the insulin secretion dose-response (starting at 10% of maximum effect), assuming similar potencies between the two effects.

*Step 4 of M4 drug discovery: Predict human exenatide treatment outcomes using all the previous steps*

The simulations of the 30-week exenatide treatment in humans (14) required five types of information to be declared.

- The first information declared was to specify the total body weight of a human in the simulation, which was set to the average body weight of the participants in the study (102 kg).
- The second information declared was to assign a basal plasma glucose concentration before exenatide treatment. As described in the main text, this was done by calibrating an initial insulin resistance to fit the first data point in the study (**Fig. 6A**, t=0; marked ‘x’).
- The third information declared determined the meal size for the repeated mixed meals. As described in the main text, this was necessary because we could only find the macronutrient profile of the meal in the human study (55% carbohydrates, 15% protein, and 30% fat) and not the meal size. The meal size calibration was achieved by calibrating a parameter that scaled the meal according to the specified macronutrient profile. In practice, this was performed to the first mixed meal consumed prior treatment (**Fig. 6A ii**, red). The calibration resulted in a 900-kcal meal.
- The fourth information declared determined the rate of insulin resistance regression rate progression to fit the initial datapoint of the second mixed meal (**Fig. 6A,** yellow; t=0; marked ‘x’). As described in the main text, this was necessary because no other data could be found that could determine the insulin resistance regression rate.
- The fifth and final information declared was the absorption rate of the long-acting release formulation. In practice, this was performed by calibrating maximum adsorption rate (Ka) and saturation (Ka_50_) to the exposure data (**Fig. 6A**, marked ‘x’). This was necessary because the slow-release formulation of exenatide (administration weekly) used in the validation study was not present in the estimation data (**Fig. 2B**). All other pharmacokinetic parameters were scaled according to the established human pharmacokinetic translation (**Fig. 2**).

All parameter calibrations listed above were done to the parameter-set corresponding to the best agreement to estimation data (**Figs. 5-6**, lines). The same calibration parameters were used across all simulations and simulation alternatives. Therefore, any differences between the simulation alternatives (**Fig. 5B,** Alt. 1-7) are caused by the different implementation of the preclinical complementary insights, generated from all previous M4 drug discovery steps (steps 1-3).

In our analysis of the 30-week human study one data denormalization step was required (**Fig. 6A i**). This was necessary because of two reasons: a) the basal plasma glucose was reported in the original publication (14) as a group relative change (mean ± SD) to participant initial value, and b) the human participants had different initial fasting plasma glucose levels 9.6 ± 2.4 mM (mean ± SD). With no reported subject-specific data we could therefore not determine basal plasma glucose concentrations without a denormalization. The denormalization was performed in order to change the unit from relative change (as reported in the data source) to basal plasma glucose (**Fig. 6A i**). This was performed by using the reported basal plasma glucose concentrations from the mixed meals first datapoints (**Fig. 6A b**, red and yellow; t=0; marked ’x’), with the assumption that the datapoint 15 minutes before the mixed meal is the basal plasma glucose. The resulting denormalization is included in the supplementary material (**Supplementary Data S8**).

The diet simulated during the 30-week exenatide treatment was assumed to consist of three standardised mixed meals every day. As described in the main text, these meals were assumed to be consumed over 15 minutes each day at 07:00, 13:00, and 18:00 o’clock. The simulation of the standardised diet began 14 days prior to the study’s start to ensure the model reached a steady state prior to the simulated study. The macronutrient profiles were assumed to follow the mixed meals reported in the article (55% carbohydrates, 15% protein, and 30% fat), and the size of the meal was assumed to be 900 kcal (2700 kcal/day), in accordance with the meal size calibration. The simulated diet was allowed to change throughout the 30-week simulation, in two different ways. The first way is through an active diet change, which the participants may have started in the beginning of the study. The active diet change was assumed to be 85% of the initial intake (2295 kcal/day) and was simulated to start at the beginning of the study in two simulation alternatives (**Fig. 5B,** Alts. 2 and 7). The exenatide-induced appetite reduction from rats and the active dietary changes were not assumed to be additive, i.e. if both diet change events occurred in the simulation the diet was simulated with 1890 kcal/day (85% of the initial intake), in accordance with the exenatide-induced reduction in appetite identified from the rat study. In accordance with the original study, the measured repeated mixed meals (**Fig 7A b,** red and yellow) were always simulated as the same meal-size (900 kcal each), independent of appetite and diet changes. This can be seen in the simulation of hepatic insulin response and hepatic glycogen (**Fig. 6B iv**), where the responses are larger at these time-points (0,14, and 29 weeks). The measured repeated mixed meal was assumed to be breakfast.

With all step-specific parts of M4 drug discovery explained the next section explains all parts of the mathematical modelling.

**Mathematical model structure**

First the pharmacokinetic model is explained and then the M4-model.

*The pharmacokinetic model*
The pharmacokinetic model was first introduced in step 2 of the analysis and was used in all subsequent steps. The model uses two states to describe exenatide pharmacokinetics from cell- and animal-studies.

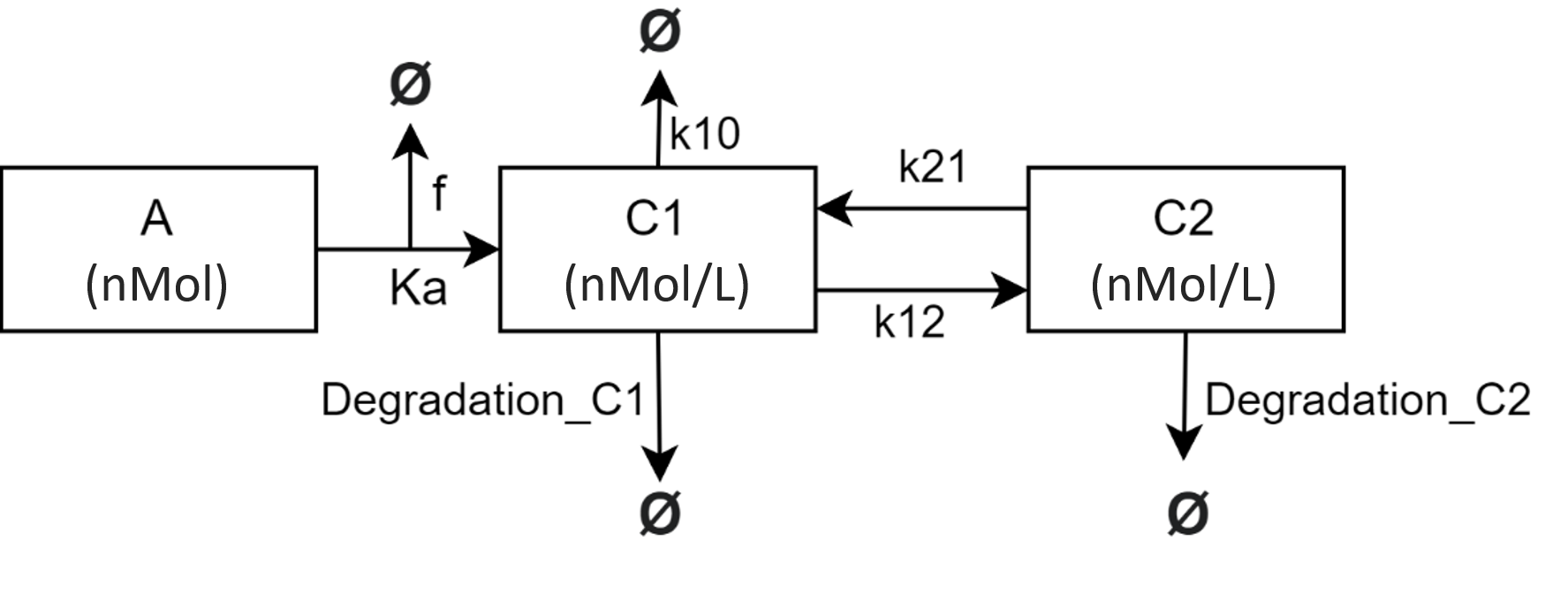

***Illustration of the pharmacokinetic model.*** *Rectangles represent states, text with no squares represents model reactions, and arrows represent metabolic rates. Flows into Ø represent flows leading out of model.*

The pharmacokinetic model includes *in vitro* clearance (Degradation_C1 and Degradation_C2), an additional *in vivo* clearance (k10), exenatide appearance from subcutaneous injections (Ka), bioavailability of exenatide administered from subcutaneous injections (f), and distribution of exenatide between a central and a peripheral compartment (k21 and k12).

##### Below are all parts of the pharmacokinetic model declared and explained: a) model states, b) model parameters, c) Model ordinary differential equations (ODEs) and reactions

##### a) Model states

The model states are time-dependent variables which are defined by ordinary differential equations (ODEs).

***List of model states***

| Name | Unit | Description | Initial value |
| --- | --- | --- | --- |
| A | nMol | Subcutaneous injections administration compartment | 0 |
| C1 | nMol/L | Exenatide in central compartment | 0 |
| C2 | nMol/L | Exenatide in non-central compartment | 0 |

##### b) Model parameters

As described above and in main text, all model parameters can be described as either base parameters (which describes general metabolic reactions), and condition alterations (which describes steps from the base parameters to specific conditions).

***List of base parameters in the pharmacokinetic model***
*θ denotes best agreement to data*

| **Parameter** | **Unit** | **Description** | **Θ** |
| --- | --- | --- | --- |
| Ka | 1/min | Maximum transportation from subcutaneous injections into central compartment | 0,597 |
| Ka50 | nMol | Amount of exenatide substance needed to achieve half of the maximum transportation rate from subcutaneous injections administration into central compartment | 46,651 |
| Q_ref | L/min | Transportation rate between state C1 and C2 | 0,001 |
| V1_ref | L | Volume of central compartment | 0,044 |
| V2_ref | L | Volume of non-central compartment | 0,052 |
| CL_ref | L/min | Maximum additional *in vivo* clearance | 1,548 |
| CL50 | nMol/L | Exenatide exposure where half the maximum additional *in vivo* clearance is reached | 348,482 |
| b_reaction | dimensionless | Scaling constant for metabolic rates based on species total body weight | 0,600 |
| b_volume | dimensionless | Scaling constant for volumes based on species total body weight | 0,997 |
| degrad | 1/min | *In vitro* clearance | 0,001 |

***List of condition alterations between species.***
*The parameter alterations influence the base parameter through multiplication, where 1 means no change and 2 means a 2-fold increase. The name of the condition alteration is identical to the base parameter which it affects, i.e. Ka in the table affects the base parameter, Ka.*

| **Condition alteration** | | **f** | **Ka** | **CL_ref** | **V1_ref** | **V2_ref** |
| --- | --- | --- | --- | --- | --- | --- |
| Description | | Bioavailability of subcutaneous injections | Maximum transportation from subcutaneous injections | Maximum additional *in vivo* clearance | Volume of central compartment | Volume of non-central compartment |
| Unit | | Step from basal parameter (fold-increase) | | | | |
| Animal species | Rat | 0,663 | 0,566 | 1,495 | 1,066 | 1,058 |
|  | Monkey | 1 | 1,736 | 0,533 | 0,652 | 1,76 |
|  | Dog | No data (fixed to 1) | | 1,661 | 1,496 | 0,585 |

The ODEs and reactions of the pharmacokinetic model are listed below.

##### c) Model ordinary differential equations (ODEs) and reactions

Exenatide administrated from subcutaneous injections were described with one ODE (**Eq. S1**).

$\frac{d}{\mathrm{dt}}\left( A \right)=- \frac{Ka*A}{Ka50+A}$	$A\left( 0 \right)=dose*f$	(Eq. S1)

where A describes an exenatide compartment representing the subcutaneous injection, and where the components of the right-hand side are defined as:
Ka denotes maximum transportation rate,
Ka50 denotes saturation of the transportation rate,
$dose$ denotes the exenatide subcutaneous injection dose, and
$f$ denotes subcutaneous injection bioavailability. Note that Ka and $f$ are described with condition alterations between animal species.

The release of exenatide from subcutaneous injections appears in the central compartment, is described with one ODE (**Eq. S2**).

$\frac{d}{\mathrm{dt}}\left( C1 \right)=\left( IVInfusion+\frac{Ka*A}{Ka50+A} \right)* \frac{1}{V1}+\left( k21*C2 \right)*\frac{V2}{V1}-(k12*C1+k10+degrad*C1)$	(Eq. S2)

where $C1$describes exenatide concentration in central compartment, and where the components of the right-hand side are defined as:
IVInfusion denotes flow of exenatide from IV infusions,
$\frac{Ka*A}{Ka50+A}$ denotes the flow of exenatide from subcutaneous injections (**Eq. S1**),
k21 denotes exenatide flow from non-central compartment (C2) to central compartment (C1), $k12$ denotes exenatide transportation from central compartment (C1) to non-central compartment (C2),
$k10$ denotes additional *in vivo* clearance,
$degrad$ denotes *in vitro* clearance,
$V1$denotes volume of central compartment ($C1$), and
$V2$denotes volume of non-central compartment ($C2)$. Note that parameters $V1$, $V2$, and $Ka$ are described with condition alterations between animal species.

The model includes a non-central compartment which is described with one ODE (**Eq. S3**).

$\frac{d}{\mathrm{dt}}\left( C2 \right)=\left( k12*C1 \right)*\frac{V1}{V2}-k21*C2 -degrad*C2$	(Eq. S3)
where $C2$describes exenatide concentration in non-central compartment, and where the components of the right-hand side are defined as:

k21 denotes exenatide transportation from non-central compartment (C2) to central compartment (C1), $k12$ denotes exenatide transportation from central compartment (C1) to non-central compartment (C2),
$degrad$ denotes *in vitro* clearance,

$V1$denotes volume of central compartment ($C1$), and
$V2$denotes volume of non-central compartment ($C2)$. Note that parameters $V1$, and $V2$, are described with condition alterations between animal species.

The exenatide rates between the central and non-central compartment are described with two equations (**Eqs. S4-S5**).

$k21=Q* \frac{1}{V2}$	(Eq. S4)
where $k21$ describes exenatide flow from non-central to central compartment, and where the components of the right-hand side are defined as:

$Q$ denotes rate between the compartments, and
$V2$denotes volume of non-central compartment ($C2)$. Note that parameter $V2$ is described with condition alterations between animal species.

$k12=Q* \frac{1}{V1}$	(Eq. S5)
where $k12$ describes exenatide flow from central to non-central compartment, and where the components of the right-hand side are defined as:

$Q$ denotes transportation rate between the compartments, and
$V1$denotes volume of central compartment ($C1)$. Note that parameter $V1$ is described with condition alterations between animal species.

The additional *in vivo* clearance is described with a dependency to exenatide in the central compartment (**Eq. S6**).

$k10=\frac{CL*C1}{CL50+C1}*\frac{1}{V1}$	(Eq. S6)

where $k10$describes additional *in vivo* clearance, and where the components of the right-hand side are defined as:
$CL$ denotes maximum additional *in vivo* clearance,
$C1$ denotes exenatide in central compartment,
$CL50$ denotes exenatide exposure of half maximum additional *in vivo* clearance, and
$V1$ denotes volume of compartment C1. Note that parameters $CL$ and $V1$ are described with condition alterations between animal species.

In the pharmacokinetic model, metabolic rates and volumes are scaled to animal total body weight. The scaling is normalised to the animal species with the lowest body weight in the estimation data, which is rats with a total body weight of 0.34 kg in the pre-existing study (1).

Two volumes are scaled to species total body weight (**Eqs. S7-8**).

$V1=V1_{ref}* {(\frac{{BW}_{act}}{{BW}_{ref}})}^{b_{volume}}$	(Eq. S7)
where $V1$ describes volume of central compartment after scaling, and where the components of the right-hand side are defined as:
$V1_{ref}$ denotes the reference volume of the central compartment before scaling,
${BW}_{act}$ denotes body weight of animal simulated,
${BW}_{ref}$ denotes body weight of reference animal, and
$b_{volume}$ denotes allometric scaling of volumes.

$V2=V2_{ref}* {(\frac{{BW}_{act}}{{BW}_{ref}})}^{b_{volume}}$	(Eq. S8)

where $V2$ describes volume of non-central compartment after scaling, and where the components of the right-hand side are defined as:
$V2_{ref}$ denotes the reference volume of the non-central compartment before scaling, ${BW}_{act}$ denotes body weight of animal simulated, ${BW}_{ref}$ denotes body weight of reference animal, and
$b_{volume}$ denotes allometric scaling of volumes.

Two metabolic rates were scaled with species total body weight (**Eqs. S9-10**).

$CL={CL}_{ref}* {(\frac{{BW}_{act}}{{BW}_{ref}})}^{b_{reaction}}$	(Eq. S9)

where $CL$ describes maximum additional *in vivo* clearance and where the components of the right-hand side are defined as:
$CL_{ref}$ denotes the reference maximum additional *in vivo* clearance,
${BW}_{act}$ denotes body weight of animal simulated,
${BW}_{ref}$ denotes body weight of reference animal, and
$b_{reaction}$ denotes allometric scaling of metabolic reactions.

$Q=Q_{ref}* {(\frac{{BW}_{act}}{{BW}_{ref}})}^{b_{reaction}}$	(Eq. S10)

where $Q$ describes transportation rate between the two pharmacokinetic compartments, and where the components of the right-hand side are defined as:
$Q_{ref}$ denotes transportation rate before scaling,
${BW}_{act}$ denotes body weight of animal simulated,
${BW}_{ref}$ denotes body weight of reference animal, and
$b_{reaction}$ denotes allometric scaling of metabolic reactions.

The pharmacokinetic model described above was integrated into the M4-model (described below).

*The M4-model*

In this section all parts of the M4-model are explained.

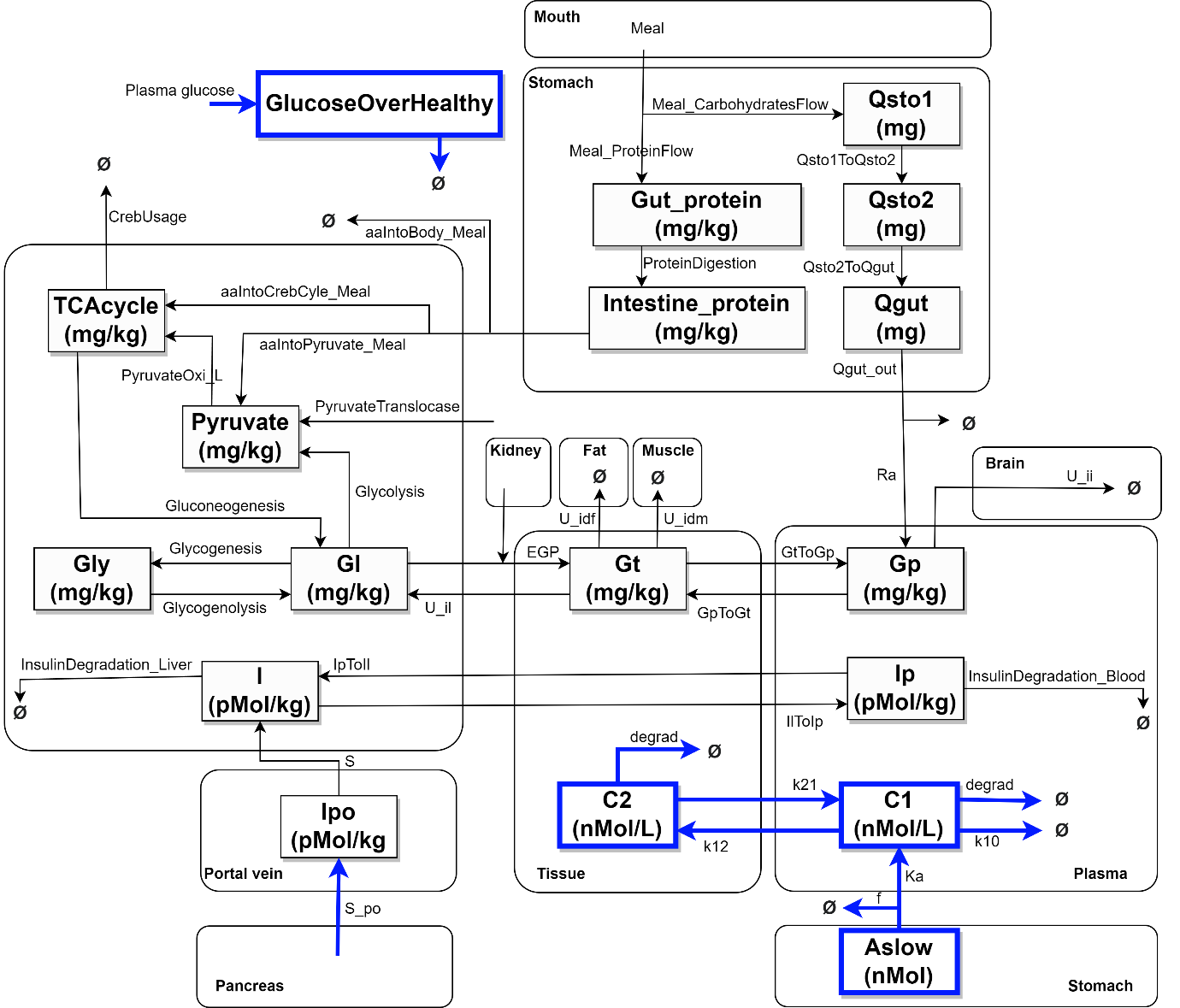
***Illustration of the M4-model***
*Rectangles represent states, text with no squares represents model reactions, and arrows represents flows. Flows into Ø are flows leading out of model. Blue represents new parts of the model and grey represents the pre-existing model.*

Each part of M4-model is declared and explained in five sections: a) model states, b) model parameters, c) model ordinary differential equations (ODEs), d) model reactions, and e) model variables.

##### a) Model states

The model states are time-dependent variables which are defined by ordinary differential equations (ODEs).

***List of model states***

| New or old | Name | Unit | Description | Initial value |
| --- | --- | --- | --- | --- |
| Old | Qsto1 | mg | First carbohydrate digestion state | Simulated |
|  | Qsto2 | mg | Second carbohydrate digestion state | Simulated |
|  | Qgut | mg | Third carbohydrate digestion state | Simulated |
|  | Intestine_Protein_ | mg/kg | First protein digestion state | Simulated |
|  | Intestine_aa_ | mg/kg | Second protein digestion state | Simulated |
|  | Gp | mg/kg | Glucose in plasma | Simulated |
|  | Ip | pmol/kg | Insulin in plasma | Simulated |
|  | Ipo | pmol/kg | Insulin in portal vein | Simulated |
|  | Gt | mg/kg | Glucose in tissue | Simulated |
|  | Gl | mg/kg | Glucose in liver | Simulated |
|  | Gly_L_ | mg/kg | Hepatic glycogen | Simulated |
|  | Pyruvate_L_ | mg/kg | Pyruvate in liver | Simulated |
|  | TCAcycle_L_ | mg/kg | Sum of components in hepatic TCA cycle | Simulated |
|  | Il | pmol/kg | Insulin in liver | Simulated |
|  | InsulinStabilization | pmol/kg | Part of insulin production calculations | Simulated |
| New | GlucoseOverHealthy | mM | A disease state that describes insulin resistance progression and regression relation to glucose levels. This state is the mathematical model describing the cell cultures. | 0 |
|  | A | nMol | Subcutaneous injections administration compartment. This state is from the pharmacokinetic model. | 0 |
|  | C1 | nMol/L | Exenatide in central compartment. This state is from the pharmacokinetic model. | 0 |
|  | C2 | nMol/L | Exenatide in non-central compartment. This state is from the pharmacokinetic model. | 0 |

##### b) Model parameters

All parameters in the model are listed below with a description of what they represent, what value was used for best agreement to data, and how that value was determined.

***List of model parameters***
θ *denotes best agreement to data.*

| Parameter | Unit | Description | θ | Parameter estimation |
| --- | --- | --- | --- | --- |
| Q_ref | 1/min | Exenatide rate between central and non-central compartment | 0,001 | Fixed from step 3 in the M4 drug discovery exemplification |
| V1_ref | L | Volume of central compartment | 0,044 |  |
| V2_ref | L | Volume of non-central compartment | 0,052 |  |
| CL_ref | 1/min | Rate of maximum additional *in vivo* clearance | 1,548 |  |
| CL50 | nMol/L | V_50_ of additional *in vivo* clearance | 348,482 |  |
| b_reaction | dimensionless | Scaling of metabolic reaction rates based on species total body weight | 0,600 |  |
| b_volume | dimensionless | Scaling of volumes based on species total body weight | 0,997 |  |
| degrad | 1/min | *In vitro* exenatide clearance | 0,0005 |  |
| KaSlow | 1/min | Maximum rate of appearance from administration | 0,04 | Estimated to calibration-data |
| KaSlow50 | nMol | V_50_ of the exenatide administration rate | 10000 |  |
| InsulinResistanceUpperlimit | mM | Glucose concentration threshold for triggering positive term of insulin resistance development | 6,708 |  |
| InsulinResistanceHealing | 1/min | Positive term for insulin resistance regression | 8,2*10^-6^ |  |
| ISk | dimensionless | Relation between disease state and insulin sensitivity | 1,4*10^-6^ |  |
| Vmax_ii | mg/kg/min | Insulin independent maximum glucose utilization of muscle and fat | 9,679 | Fixed from pre-existing model |
| Vmax_id | 1/min | Insulin dependent maximum glucose utilization of muscle and fat | 0,750 |  |
| U50 | mg/kg | Glucose exposure at half of maximum muscle and fat glucose utilization | 50,574 |  |
| U_ratio | % | How much of the muscle and fat glucose utilisation is from muscle | 70,000 |  |
| EGP_KidneysK | 1/min | Part of the renal glucose production calculation | 0,761 |  |
| GLUT2_diffusionMax | 1/min | Maximum glucose rate from liver into tissue | 6,357 |  |
| GLUT2_diffusion50 | mg/kg | Glucose exposure when glucose rate into tissue from liver is half of the maximum rate | 1,725 |  |
| Uidl_diffusionMax | 1/min | Maximum glucose rate into the liver from tissue | 1,03 |  |
| Uidl_diffusion50 | mg/kg | Glucose exposure when glucose rate into liver from tissue is half of the maximum rate | 1,106 |  |
| ExenatideInsulinMax | fold-increase | Positive term of exenatide dose-response curve | 2,051 | Fixed from previous M4 drug discovery steps using e.g. rat potency or human cell potency |
| ExenatideInsulin50 | nM | Exenatide EC_50_ | 0,206 |  |
| ExenatideInsulin_hill | dimensionless | Exenatide hill coefficient | 3,252 |  |
| ExenatideInsulinDecayMax | fold-increase | Negative term of exenatide dose-response curve | 0,316 |  |
| ExenatideInsulinDecay50 | nM | Exenatide IC_50_ | 25,069 |  |
| Exanitide_IP | % | Exenatide effect on Phase-II/Phase-I insulin production | 41,552 |  |
| k_gri | 1/min | Carbohydrate rate from first to second digestion state | 52,942 | Fixed from pre-existing model |
| k_min | 1/min | Minimum rate of gastric emptying | 0,000149 |  |
| k_max | 1/min | Maximum rate of gastric emptying | 0,00663 |  |
| b | dimensionless | Transition between minimum and maximum rate of gastric emptying | 69,689 |  |
| d | dimensionless | Transition between minimum and maximum rate of gastric emptying | 0,026 |  |
| k_abs | 1/min | Glucose rate out of intestines into plasma | 0,035 |  |
| ProteinBreakdown | 1/min | Protein rate from first to second digestion state | 0,0013 |  |
| aaTransportation_k | 1/min | Rate of amino acids out of intestines | 2,676 |  |
| f | ratio | Carbohydrate bioavailability | 0,810 |  |
| k_1 | 1/min | Rate of glucose from blood to tissue | 0,046 |  |
| U_ii | mg/kg/min | Insulin-independent glucose utilization | 0,766 |  |
| m_2 | 1/min | Rate of insulin from blood to liver | 0,286 |  |
| m_4 | 1/min | Non-hepatic insulin clearance | 0,0097 |  |
| K | dimensionless | Glucose rate of change conversion to insulin production | 179,958 |  |
| S_b | pmol/kg/min | Basal insulin production | 3,1*10^-5^ |  |
| gamma | 1/min | Rate of insulin secretion | 8273,37 |  |
| k_2 | 1/min | Rate of glucose from tissue to blood | 0,910 |  |
| G_b | mg/kg | Basal glucose concentration before diseased state multiplication | 30,336 |  |
| V_glyBmax | 1/min | Maximum rate of glycogenolysis | 1,180 |  |
| GlyB | mg/kg | Amount of glycogen glucose at half of maximum glycogenolysis | 1,752 |  |
| V_glySmax | 1/min | Maximum rate of glycogenesis | 1,670 |  |
| GlyS | mg/kg | Amount of hepatic glycogen at half of maximum glycogenesis | 1,983 |  |
| CrebUsage_K | 1/min | Rate of disappearance of components in the TCA cycle | 0,0016 |  |
| Gluconeogenesis_CrebK | 1/min | Gluconeogenesis from the TCA cycle in the liver | 0,486 |  |
| PyruvateTranslocase_K | 1/min | Rate of appearance of pyruvate from the body into the liver | 0,00039 |  |
| PyruvateOxi_k | 1/min | Rate of appearance of components in TCA cycle from pyruvate | 0,0020 |  |
| Aminoprofile_k | % | Ratio of amino acids which appears in the hepatic metabolism as pyruvate | 94,99 |  |
| Glycolysis_k | 1/min | Rate of hepatic glycolysis | 0,363 |  |
| m_1 | 1/min | Rate of insulin from liver to blood | 0,329 |  |
| m_5 | 1/min | Negative term in hepatic insulin extraction | 1,036 |  |
| m_6 | 1/min | Positive term in hepatic insulin extraction | 0,502 |  |
| alpha | 1/min | Delay between plasma glucose change and insulin production | 3,181 |  |
| beta | dimensionless | Scaling of insulin production based on glucose concentration | 0,0185 |  |
| GlyDep_Meal | mMol/L | Basal glycogen as a proxy for the fed state of the body when regulating amino acid meal distribution | 388,317 |  |
| GlyDep_TCA | mMol/L | Basal glycogen as a proxy for the fed state of the body when regulating TCA component utilisation | 46,992 |  |
| GlyDep_Glugoneogenesis | mMol/L | Basal glycogen as a proxy for the fed state of the body when regulating hepatic gluconeogenesis | 2657,89 |  |
| GlyDep_KidneysEGP | mMol/L | Basal glycogen as a proxy for the fed state of the body when regulating renal glucose production and flow into hepatic pyruvate from the body | 0,093 |  |
| GlyDepEXP_Meal | Dimensionless | Exponent for glycogen as a proxy for the fed state of the body when regulating amino acid meal distribution | 3,000 |  |
| GlyDepEXP_TCA | Dimensionless | Exponent for glycogen as a proxy for the fed state of the body when regulating TCA component utilisation | 1,099 |  |
| GlyDepEXP_Glugoneogenesis | Dimensionless | Exponent for glycogen as a proxy for the fed state of the body when regulating hepatic gluconeogenesis | 0,100 |  |
| GlyDepEXP_KidneysEGP | Dimensionless | Exponent for glycogen as a proxy for the fed state of the body when regulating renal glucose production and flow into hepatic pyruvate from the body | 0,134 |  |
| Meal_ProteinFlow | mg/min | Protein flow from meal | Model inputs |  |
| Meal_CarbohydratesFlow | mg/min | Carbohydrate flow from meal |  |  |
| GlucoseInjection | mg/kg/min | Glucose injection |  |  |
| D | mg | Total carbohydrate amount in meal |  |  |
| BW | kg | Total body weight |  |  |
| IVInfusion | uMol/min | Used when simulation exenatide IV infusions |  |  |

The ODEs of all model states are listed below.

##### c) Model ordinary differential equations (ODEs)

Three model states represent the digestion of carbohydrate from meals (**Eqs. S11-S13**).

$\frac{d}{\mathrm{dt}}\left( Qsto1 \right)=\left( \mathrm{CarbohydratesFlow} \right)-\left( Qsto1toQsto2 \right)$	(Eq. S11)

where $Qsto1$ describes the first of three digestive steps when consuming carbohydrates from meals, and where the components of the right-hand side are defined as:

$\mathrm{CarbohydratesFlow}$ denotes carbohydrate input from meals, and
$Qsto1toQsto2$ denotes the flow of carbohydrates from the first to the second digestive state.

$\frac{d}{\mathrm{dt}}\left( Qsto2 \right)=\left( Qsto1toQsto2 \right)-\left( Qsto2ToQgut \right)$	(Eq. S12)
where $Qsto2$ describes the second of three digestive steps when consuming carbohydrates from meals, and where the components of the right-hand side are defined as:

$Qsto1toQsto2$ denotes a flow from the first to the second digestive state, and
$Qsto2ToQgut$ denotes a flow from the second to the third and final digestive state.

$\frac{d}{\mathrm{dt}}\left( \mathrm{Qgut} \right)=\left( Qsto2ToQgut \right)-\left( \mathrm{Qgu}t_{\mathrm{out}} \right)$	(Eq. S13)

where $\mathrm{Qgut}$ describes the third of three digestive steps when consuming carbohydrates from meals, and where the components of the right-hand side are defined as:

Where $Qsto2ToQgut$ denotes flow from the second to the third and final digestive state, and $\mathrm{Qgu}t_{\mathrm{out}}$ denotes the output of the carbohydrate digestion system.

Two model states describe protein digestion from meals (**Eqs. S14-S15**).

$\frac{d}{\mathrm{dt}}\left( \mathrm{Intestines}_{\mathrm{Protein}} \right)=\left( \frac{\mathrm{Protei}n_{\mathrm{Meal}}}{\mathrm{BW}} \right)-\left( \mathrm{ProteinDigestion} \right)$	(Eq. S14)

where $\mathrm{Intestines}_{\mathrm{Protein}}$ describes the first of two digestive steps when consuming proteins from meals, and where the components of the right-hand side are defined as:

$\mathrm{Protei}n_{\mathrm{Meal}}$ denotes protein input from meals,
$\mathrm{BW}$ denotes species total body weight, and
$\mathrm{ProteinDigestion}$ denotes protein digestion from the first to the second digestion state.

$\frac{d}{\mathrm{dt}}\left( \mathrm{Intestines}_{\mathrm{aa}} \right)=ProteinDigestion -\left( \mathrm{aaTransportatio}n_{\mathrm{Meal}} \right)$	(Eq. S15)

where $\mathrm{Intestines}_{\mathrm{aa}}$ describes the second of two digestive steps when consuming proteins from meals, and where the components of the right-hand side are defined as:

$\mathrm{ProteinDigestion}$ denotes protein digestion from the first to the second digestion state, and
$\mathrm{aaTransportatio}n_{\mathrm{Meal}}$ denotes output of the protein digestion system.

Three model states describe glucose in different compartments of the body (**Eqs. S16-S18**).

$\frac{d}{\mathrm{dt}}\left( \mathrm{Gp} \right) =\left( Ra+ GtToGp \right)-\left( U_{\mathrm{ii}}+ GpToGt \right)$	(Eq. S16)
where $\mathrm{Gp}$ describes plasma glucose, and where the components of the right-hand side are defined as:

$\mathrm{Ra}$ denotes plasma glucose rate of appearance from the carbohydrate digestion system,
$\mathrm{GtToGp}$ denotes glucose transportation rate from tissue to plasma,
$U_{\mathrm{ii}}$ denotes brain insulin independent glucose utilisation, and
$\mathrm{GpToGt}$ denotes glucose transportation rate from plasma to tissue.

$\frac{d}{\mathrm{dt}}\left( \mathrm{Gt} \right)=\left( GlucoseInjection+ GpToGt+EGP \right)-\left( U_{\mathrm{idf}}+ U_{\mathrm{idm}}+ GtToGp+ U_{\mathrm{il}} \right)$	(Eq. S17)
where $\mathrm{Gt}$ describes glucose in tissue, and where the components of the right-hand side are defined as:

$\mathrm{GlucoseInjection}$ denotes non intravenous glucose injections,
$\mathrm{GpToGt}$ denotes glucose transportation rate from plasma to tissue,
$\mathrm{EGP}$ denotes glucose transportation from kidneys and liver to tissue,
$U_{\mathrm{idf}}$denotes fat glucose utilisation,
$U_{\mathrm{idm}}$denotes muscle glucose utilisation,
$\mathrm{GtToGp}$ denotes glucose transportation rate from tissue to plasma, and
$U_{\mathrm{iil}}$ denotes glucose flow from tissue to liver.

$\frac{d}{\mathrm{dt}}\left( \mathrm{Gl} \right)=\left( U_{\mathrm{il}}+ Glycogenolysis+Gluconeogenesis \right)-\left( Glycogenesis+ EGP_{\mathrm{Liver}}+Glycolysis \right)$	(Eq. S18)
where $\mathrm{Gl}$ describes glucose in hepatic glucose, and where the components of the right-hand side are defined as:

$U_{\mathrm{iil}}$ denotes glucose flow from tissue to liver,
$\mathrm{Glycogenolysis}$ denotes hepatic glycogenolysis,
$\mathrm{Gluconeogenesis}$denotes hepatic gluconeogenesis,
$\mathrm{Glycogenesis}$ denotes hepatic glycogenesis,
$\mathrm{EG}P_{\mathrm{Liver}}$ denotes glucose flow out of liver to tissue, and
$\mathrm{Glycolysis}$ denotes hepatic glycolysis.

Three states describe hepatic glycogen, pyruvate, and a sum of metabolites in the hepatic TCA cycle (**Eqs. S19-S21**)

$\frac{d}{\mathrm{dt}}\left( \mathrm{Gl}y_{L} \right)=\left( \mathrm{Glycogenesis} \right)-\left( \mathrm{Glycogenolysis} \right)$	(Eq. S19)
where $\mathrm{Gl}y_{L}$describes hepatic glycogen, and where the components of the right-hand side are defined as:

$\mathrm{Glycogenesis}$ denotes hepatic glycogenesis, and
$\mathrm{Glycogenolysis}$ denotes hepatic glycogenolysis.

$\frac{d}{\mathrm{dt}}\left( \mathrm{Pyruvat}e_{L} \right)= \left( \mathrm{aaIntoPyruvat}e_{\mathrm{Meal}}+Glycolysis+PyruvateTranslocase \right)-\left( \mathrm{PyruvateOxi}_{L} \right)$	(Eq. S20)
where $\mathrm{Pyruvat}e_{L}$ describes hepatic pyruvate, and where the components of the right-hand side are defined as:
$\mathrm{aaIntoPyruvat}e_{\mathrm{Meal}}$ denotes hepatic pyruvate rate of appearance from meals,
$\mathrm{Glycolysis}$ denotes hepatic glycolysis, $\mathrm{PyruvateTranslocase}$ denotes a flow from the body into hepatic pyruvate, and
$\mathrm{PyruvateOxi}_{L}$ denotes hepatic pyruvate utilisation.

$\frac{d}{\mathrm{dt}}\left( \mathrm{TCAcycl}e_{L} \right)=\left( \mathrm{aaIntoTCAcycl}e_{\mathrm{Meal}}+PyruvateOxi_{L} \right)-\left( \mathrm{Gluconeogenesi}s+TCA_{\mathrm{usage}} \right)$	(Eq. S21)
where $\mathrm{TCAcycl}e_{L}$describes a summarised and simplified state of hepatic TCA cycle components, and where the components of the right-hand side are defined as:
$\mathrm{aaIntoTCAcycl}e_{\mathrm{Meal}}$ denotes hepatic TCA metabolites appearance from meals, $\mathrm{PyruvateOx}i_{L}$denotes hepatic TCA metabolites metabolised from hepatic pyruvate, $\mathrm{Gluconeogenesi}s$ denotes hepatic gluconeogenesis, and
$\mathrm{TC}A_{\mathrm{usage}}$ denotes a loss of metabolites in the hepatic TCA cycle out of the model.

Three model states describe insulin in various compartments of the body (**Eqs. S22-S24**)

$\frac{d}{\mathrm{dt}}\left( \mathrm{Ipo} \right)=Spo-S$	(Eq. S22)
where $\mathrm{Ipo}$ describes insulin in the portal vein, and where the components of the right-hand side are defined as:

$\mathrm{Spo}$ denotes insulin production, and
$S$ denotes insulin secretion.

$\frac{d}{\mathrm{dt}}\left( \mathrm{Il} \right)=\left( S+ IpToIl \right)-\left( IlToIp+ \mathrm{InsulinDegradation}_{\mathrm{Liver}} \right)$	(Eq. S23)
where $\mathrm{Il}$ denotes hepatic insulin, and where the components of the right-hand side are defined as:

$S$ denotes insulin secretion,
$\mathrm{IpToIl}$ denotes insulin transportation from plasma to liver,
$\mathrm{IlToIp}$ denotes insulin transportation from liver to plasma, and
$\mathrm{InsulinDegradation}_{\mathrm{Liver}}$ denotes hepatic insulin clearance.

$\frac{d}{\mathrm{dt}}\left( \mathrm{Ip} \right)=\left( \mathrm{IlToIp} \right)-\left( IpToIl+ \mathrm{InsulinDegradation}_{\mathrm{Blood}} \right)$	(Eq. S24)
where $\mathrm{Ip}$ describes plasma insulin, and where the components of the right-hand side are defined as:

$\mathrm{IlToIp}$ denotes insulin transportation from liver to plasma,
$\mathrm{IpToIl}$ denotes insulin transportation from plasma to liver, and $\mathrm{InsulinDegradation}_{\mathrm{Blood}}$ denotes non-hepatic insulin clearance.

One model state is used to describe insulin production dependent on glucose levels (**Eq. S25**)

$\frac{d}{\mathrm{dt}}(InsulinStabilization) =\frac{\mathrm{alpha}}{180.16}*\left( beta*\left( Gp-{IS*G}_{b} \right)-\mathrm{InsulinStabilization} \right)$	(Eq. S25)
where $\mathrm{InsulinStabilization}$ describes plasma glucose concentration dependent insulin production, which is based on:

$\mathrm{alpha}$ denotes a delay rate between insulin production and glucose concentrations,
180.16 denotes glucose molecular weight,
$\mathrm{beta}$ denotes magnitude of insulin produced,
$\mathrm{IS}$ denotes the new insulin sensitivity function implemented from the human cell cultures which moves the basal plasma glucose threshold in accordance with disease progression state ($\mathrm{GlucoseOverHealthy}$**; Eq. S26**), and
$G_{b}$ denotes plasma glucose concentration triggering threshold based on disease state.

One new model state describes insulin resistance progression and regression (**Eq. S26**).

$\frac{d}{\mathrm{dt}}(GlucoseOverHealthy) =({\max\left( 0, \frac{G}{18}-InsulinResistanceUpperlimit \right))}^{2}-InsulinResistanceHealing*GlucoseOverHealthy$	(Eq. S26)
where $\mathrm{GlucoseOverHealthy}$ describes long-term changes to insulin sensitivity from short-term glucose concentrations changes from meals, and where the components of the right-hand side are defined as:

$\frac{G}{18}$ denotes plasma glucose (mM),
$InsulinResistanceUpperlimit$ denotes a threshold when glucose concentrations increase diseased state, and
$InsulinResistanceHealing$ denotes rate of insulin resistance regression.

The final ODEs used in the M4-model is the exenatide pharmacokinetic model, which was described above (**Eqs. S1-S3**).

All model states described above are defined by metabolic reactions, which are declared below.

##### d) Model metabolic reactions

The metabolic reactions used in the pharmacokinetic model are also used in the M4-model to simulate exenatide pharmacokinetics (**Eqs. S4-S6**).

Changes to insulin sensitivity is described with a dependency to the new disease state (**Eq. S27**).

$IS=1+\mathrm{ISk}* GlucoseOverHealthy$	(Eq. S27)
where $\mathrm{IS}$ describes insulin sensitivity changes, and where the components of the right-hand side are defined as:

1 represents the baseline level in the absence of the diseased state,
$\mathrm{ISk}$ denotes the insulin sensitivity relation to the diseased state, and
$\mathrm{GlucoseOverHealthy}$ detonate the diseased state.

Exenatide effect on insulin production is imported from the mathematical modelling of exenatide effect on human cell cultures. The effect production was described with a dependency on exenatide in the media (implemented as plasma) (**Eq. S28**).

$\mathrm{Exenatide}_{\mathrm{PD}}= \frac{ExenatideInsulinMax * {Exenatide}_{Medium}^{Exenatide_{Hill}}}{{ExenatideInsulin50}^{Exenatide_{Hill}} + {Exenatide}_{Medium}^{Exenatide_{Hill}}}- \frac{ExenatideInsulinDecayMax * {Exenatide}_{Medium}^{Exenatide_{Hill}}}{{ExenatideInsulinDecay50}^{Exenatide_{Hill}} + {Exenatide}_{Medium}^{Exenatide_{Hill}}}$	(**Eq. S28**)

where $\mathrm{Exenatide}_{\mathrm{PD}}$ describes the bell-shaped exenatide dose-response effect on insulin secretion and production, and where the components of the right-hand side are defined as:
$ExenatideInsulinMax$ and $ExenatideInsulinDecayMax$ denote the maximum effect of both the positive and negative parts of the bell-shaped dose-response curve, $Exenatide_{Hill}$ denotes hill coefficient from no response to maximum response,
${Exenatide}_{Medium}$ denotes exenatide concentration in the media, $ExenatideInsulin50$ and $ExenatideInsulinDecay50$ denote EC_50_ and IC_50_ of the bell-shaped dose-response curve. The implementation of human donor and cell line batch variability was applied to $ExenatideInsulin50$ and $ExenatideInsulinDecay50$ (1 parameter), and to $ExenatideInsulinMax$, and $ExenatideInsulinDecayMax$ (1 parameter).

Hepatic insulin response was also imported from the description of exenatide effect on human cell cultures. In the M4-model hepatic insulin response is called “$InsulinResponseNew$” to signify the change from the previous equation called “$InsulinResponse"$. A key difference between the old and new equation is that $InsulinResponse$ had a linear relation to hepatic insulin, while the new $InsulinResponseNew$ is described with a Michaelis Menten equation (**Eq. 29**). In practice, this means that the new equation will have a maximum insulin response. A maximum hepatic insulin response was observed human cell culture experiments, and the same parameter was also used to the *in vivo* human studies ($InsulinResponseMax)$.

$InsulinResponse = \frac{\mathrm{InsulinResponseMax}* {Insulin}_{Medium}^{InsulinResponse_{Hill}}}{{(IS* InsulinResponse50)}^{InsulinResponse_{Hill}} + {Insulin}_{Medium}^{InsulinResponse_{Hill}}}$	(**Eq. S29**)

where $InsulinResponse$ describes an increase in insulin dependent reaction rates, and where the components of the right-hand side are defined as:
$\mathrm{InsulinResponseMax}$ denotes maximum response amplitude,
$Insulin_{Medium}$ denotes insulin concentration in media, $InsulinResponse_{Hill}$ denotes hill coefficient of the change from no response to maximum response,
$InsulinResponse50$ denotes insulin exposure when $InsulinResponse$ reaches half of maximum effect, and $IS$ denotes insulin resistance progression in form of a factor greater or equal to 1 that reduces the potency of the insulin response.

The carbohydrate digestive system (**Eqs. S30-S32**) is kept identical to the pre-existing model (4), which in return is a further development of the ‘Nyman model’ (15) based on the original ‘Dalla Man model’ (16). For a more detailed description of these equations, we refer to the source of the equations —the original ‘Dalla Man model’. In brief, the rate of digestion is dependent on the total amount of carbohydrates in the states $Q_{sto1}$ and $Q_{sto2}$ (**Eq. S30**).

$Q_{\mathrm{sto}}=Q_{sto1}+Q_{sto2}$	(Eq. S30)

where $Q_{\mathrm{sto}}$ describes the sum of the first two carbohydrate digestive system states, and where the components of the right-hand side are defined as:

$Q_{sto1}$ denotes polysaccharides in the digestive system, and
$Q_{sto2}$ denotes monosaccharides in the digestive system.

The release of glucose from the carbohydrate digestive system is dependent on the size of the meal (**Eqs. S31-S32**).

$aa= \frac{2.5*D}{1-b}$	(Eq. S31)

where $\mathrm{aa}$ is a data-driven equation set to describe the gastric emptying from meals, and where the components of the right-hand side are defined as:

2.5 denotes a systemic constant fixed from the original publication,
$D$ denotes the total amount of carbohydrate in the latest meal, and
b denotes a parameter which creates a delay of minimum rate of gastric emptying.

$cc= \frac{2.5*D}{d}$	(Eq. S32)
where $\mathrm{cc}$ is a data-driven equation set to describe the gastric emptying from meals, and where the components of the right-hand side are defined as:

2.5 is a systemic constant fixed from the original publication,
$D$ denotes the total amount of carbohydrate in the latest meal, and
d denotes a parameter which creates a delay of maximum rate of gastric emptying.

The two data-driven gastric emptying functions (**Eqs. S31-S32**) are used in the function $K_{\mathrm{empt}}$, which is kept identical to the original publication (**Eq. S33**).

$K_{\mathrm{empt}}=K_{\min}+\frac{K_{\max}-K_{\min}}{2}*\tanh\left( aa*\left( Q_{\mathrm{sto}}-b*D \right) \right)-\tanh\left( cc*\left( Q_{\mathrm{sto}}-d*D \right) \right)+2)$	(Eq. S33)

where $K_{\mathrm{empt}}$ describes the gastric emptying of carbohydrates from meals, and where the components of the right-hand side are defined as:

$K_{\max}$ denotes maximum digestion rate,
$K_{\min}$ denotes minimum digestion rates,
$Q_{\mathrm{sto}}$ denotes total carbohydrates in the digestive system,
D denotes the total carbohydrates in the consumed meal, and
$\mathrm{aa}$ and $\mathrm{cc}$ are the data-driven equations set to describe the gastric emptying from meals (**Eqs. S31-S32**).

The transportation rates between the carbohydrate digestive system states are described identically to the pre-existing model (**Eq. S34-S36**) (4).

$Qsto1ToQsto2 =K_{\mathrm{gri}}*Qsto1$	(Eq. S34)

where $Qsto1ToQsto2$ describes the carbohydrate flow from the first to the second carbohydrate digestive state, and where the components of the right-hand side are defined as:

$K_{\mathrm{gri}}$ denotes the transportation rate, and
$Qsto1$ denotes the carbohydrates in the first carbohydrate digestive state.

${Qsto2ToQgut =K}_{\mathrm{empt}} *Qsto2$	(Eq. S35)
where $Qsto2ToQgut$ describes the carbohydrate flow from the second to the third and final carbohydrate digestive state, and where the components of the right-hand side are defined as:

$K_{\mathrm{empt}}$ denotes the transportation rate, and
$Qsto2$ denotes the carbohydrates in the second carbohydrate digestive state.

$\mathrm{Qgu}t_{\mathrm{out}}=K_{\mathrm{abs}}*Qgut$	(Eq. S36)

where $\mathrm{Qgu}t_{\mathrm{out}}$ describes the carbohydrate flow out of the final carbohydrate digestive state, and where the components of the right-hand side are defined as:

$K_{\mathrm{abs}}$ denotes the transportation rate, and
$\mathrm{Qgut}$ denotes the carbohydrates in the third carbohydrate digestive state.

The simplified protein digestive system consists of a breakdown of protein into amino acids (**Eq. S37**), and transportation rates which distributes the amino acids in the body (**Eqs. S38-S39**).

$ProteinDigestion=ProteinBreakdown * \mathrm{Intestines}_{\mathrm{Protein}}$	(Eq. S37)
where $\mathrm{ProteinDigestion}$ describes protein breakdown to amino acids, and where the components of the right-hand side are defined as:

$\mathrm{ProteinBreakdown}$ denotes the reaction rate, and
$\mathrm{Intestines}_{\mathrm{Protein}}$ denotes amount of protein in the digestion state.

$\mathrm{aaInto}\mathrm{TCAcycle}_{\mathrm{Meal}}=\left( 1-Aminoprofile_{K} \right) * ProteinMeal_{K} * \mathrm{aaTransportation}_{k}*GlyDep_{\mathrm{MealNegative}}$	(Eq. S38)
where $\mathrm{aaInto}\mathrm{TCAcycle}_{\mathrm{Meal}}$ describes transportation rate into the hepatic TCA cycle, and where the components of the right-hand side are defined as:

$\mathrm{Aminoprofil}e_{K}$ denotes the ratio of how much amino acids from the meal becomes pyruvate in the liver,
$\mathrm{ProteinMea}l_{K}$ denotes the transportation rate,
$\mathrm{aaTransportation}_{k}$ denotes amino acids distribution rate to various compartments of the body, $\mathrm{GlyDe}p_{\mathrm{MealNegative}}$ denotes a transportation rate dependency on the fed state of the body. This fed state of the body is described with a relation to hepatic glycogen. The usage of hepatic glycogen as a proxy for the fed state of the body is used in several equations of the original model to describe a simplified glucose metabolism. In practice, in this specific $\mathrm{aaInto}\mathrm{TCAcycle}_{\mathrm{Meal}}$ function, the dependency to the fed state of the body means that if protein is consumed in a fasted state more protein will be transported into the liver where gluconeogenesis is upregulated.

$\mathrm{aaInto}\mathrm{Pyruvate}_{\mathrm{Meal}} =Aminoprofile_{K} * ProteinMeal_{K}* \mathrm{aaTransportation}_{k}*GlyDep_{\mathrm{MealNegative}}$	(Eq. 39)
where $\mathrm{aaInto}\mathrm{Pyruvate}_{\mathrm{Meal}}$describes transportation of amino acids into hepatic pyruvate, and where the components of the right-hand side are defined as:

$\mathrm{Aminoprofil}e_{K}$ denotes the ratio of how much amino acids from the meal becomes pyruvate in the liver,

$\mathrm{ProteinMea}l_{K}$ denotes the transportation rate,
$\mathrm{aaTransportation}_{k}$denotes amino acids distribution to various compartments of the body, $\mathrm{GlyDe}p_{\mathrm{MealNegative}}$ denotes a transportation rate dependent on the fed state of the body.

In the model not all amino acids from meals appears in the liver. The rate of amino acids which does not appear in the liver is described with one equation regulated with a positive dependency to the fed state of the body (**Eq. S40**).

$\mathrm{aaIntoBod}y_{\mathrm{Meal}} = ProteinMeal_{K}* \mathrm{aaTransportation}_{k}**GlyDep_{\mathrm{MealPositive}}$	(Eq. S40)

where $\mathrm{aaIntoBod}y_{\mathrm{Meal}}$ describes amino acids which do not appear in the liver, and where the components of the right-hand side are defined as:

$\mathrm{ProteinMea}l_{K}$ denotes the transportation rate,
$\mathrm{aaTransportation}_{k}$ denotes amino acids which is distributed to various compartments in the body, $\mathrm{GlyDe}p_{\mathrm{MealPositive}}$ denotes a transportation rate dependent on the fed state of the body.

The three amino acid flows to various parts of the metabolism (**Eq. S38-S40**) are summarised with two equations (**Eqs. S41-S42**).

$\mathrm{aaIntoLive}r_{\mathrm{Meal}}=aaInto\mathrm{Pyruvate}_{\mathrm{Meal}} + aaInto\mathrm{TCAcycle}_{\mathrm{Meal}}$	(Eq. S41)
where $\mathrm{aaIntoLive}r_{\mathrm{Meal}}$ describes the total amino acid rate to the liver, and where the components of the right-hand side are defined as:

$\mathrm{aaInto}\mathrm{Pyruvate}_{\mathrm{Meal}}$ denotes amino acids from meals metabolised into hepatic pyruvate, and $\mathrm{aaInto}\mathrm{TCAcycle}_{\mathrm{Meal}}$ denotes amino acids from meals metabolised into components of the hepatic TCA cycle.

$\mathrm{aaTransportatio}n_{\mathrm{Meal}} = aaIntoLiver_{\mathrm{Meal}}+ aaIntoBody_{\mathrm{Meal}}$	(Eq. S42)
where $\mathrm{aaTransportatio}n_{\mathrm{Meal}}$ describes the total transportation rate of amino acids from the digestive system, and where the components of the right-hand side are defined as:

$\mathrm{aaIntoLive}r_{\mathrm{Meal}}$ denotes the rate of amino acids to the liver from meals, and
$\mathrm{aaIntoBod}y_{\mathrm{Meal}}$ denotes the rate of amino acids which are not metabolised in the hepatic metabolism.

Similarly to the protein digestive system, there is a fraction of consumed carbohydrates which is lost in digestive system and does not appear in the plasma glucose (**Eq. S43**).

$Ra =f*\frac{\mathrm{Qgu}t_{\mathrm{out}}}{\mathrm{BW}}$	(Eq. S43)
where $\mathrm{Ra}$ describes the plasma glucose rate of appearance, and where the components of the right-hand side are defined as:

$f$ denotes carbohydrate bioavailability,
$\mathrm{Qgu}t_{\mathrm{out}}$ denotes the glucose rate out of the digestive system, and
$\mathrm{BW}$ denotes subject total body weight.

The transportation of glucose between plasma to tissue is described with two equations (**Eqs. S44-S45**)

$GtToGp=K_{2}*Gt$	(Eq. S44)
where $\mathrm{GtToGp}$ describes glucose transportation from tissue to plasma, and where the components of the right-hand side are defined as:

$K_{2}$ denotes the transportation rate, and
$\mathrm{Gt}$ denotes glucose in tissue.

$GpToGt=K_{1}*Gp$	(Eq. S45)
where $\mathrm{GpToGt}$ describes glucose transportation from plasma to tissue, and where the components of the right-hand side are defined as:

$K_{1}$ denotes the transportation rate, and
$\mathrm{Gp}$ denotes glucose in plasma.

The rate of glucose from plasma to liver is described with one equation (**Eq. S46**).

$U_{\mathrm{il}}= \frac{\mathrm{Ui}\mathrm{dl}_{\mathrm{diffusionMAX}}*\mathrm{Gt}}{\mathrm{Uid}l_{diffusion50}+ Gt}$	(Eq. S46)
where $U_{\mathrm{il}}$ describes the glucose rate from tissue to liver, and where the components of the right-hand side are defined as:

$\mathrm{Ui}\mathrm{dl}_{\mathrm{diffusionMAX}}$ denotes the maximum transportation rate,
$\mathrm{Gt}$ denotes glucose in tissue, and $\mathrm{Uid}l_{diffusion50}$ denotes the glucose exposure when half of the maximum transportation rate is reached.

The glucose production is assumed to be either from renal production (**Eq. S47**) or hepatic production (**Eq. S48**). Both glucose production sources are summarised as the total endogenous glucose production (EGP) (**Eq. S49**)

$\mathrm{EG}P_{\mathrm{Kidneys}}=EGP_{\mathrm{KidneysK}}*GlyDep_{\mathrm{inflow}}$	(Eq. S47)
where $\mathrm{EG}P_{\mathrm{Kidneys}}$ describes the rate of glucose production in the kidneys, and where the components of the right-hand side are defined as:

$\mathrm{EG}P_{\mathrm{KidneysK}}$ denotes basal renal glucose production, and
$\mathrm{GlyDe}p_{\mathrm{inflow}}$ denotes a negative dependency to the fed state of the body, i.e. if hepatic glycogen is high then the kidney glucose production is downregulated.

$\mathrm{EG}P_{\mathrm{Liver}}= \frac{GLUT2diffusion_{\mathrm{MAX}}* Gl}{GLUT2diffusion_{50} + Gl}$	(Eq. S48)
where $\mathrm{EG}P_{\mathrm{Liver}}$ describes rate of glucose from the liver into tissue, and where the components of the right-hand side are defined as:

$GLUT2diffusion_{\mathrm{MAX}}$ denotes the maximum transportation rate,
$\mathrm{Gl}$ denotes glucose in the liver, and
$GLUT2diffusion_{50}$ denotes the glucose exposure when half of the maximum transportation rate is reached.

$EGP=EGP_{\mathrm{Liver}}+EGP_{\mathrm{Kidneys}}$	(Eq. S49)
where $\mathrm{EGP}$ describes the sum of endogenous glucose production, and where the components of the right-hand side are defined as:

$\mathrm{EG}P_{\mathrm{Liver}}$ denotes glucose produced by the liver, and
$\mathrm{EG}P_{\mathrm{Kidneys}}$ denotes glucose produced by the kidneys.

The muscle and fat glucose utilisation are described with four equations (**Eqs. S50-S53**).

$V_{\mathrm{maxU}}=V_{\mathrm{maxii}}+V_{\mathrm{maxid}}*\frac{I}{IS}$	(Eq. S50)
where $V_{\mathrm{maxU}}$ describes maximum glucose utilisation of both muscle and fat tissue, and where the components of the right-hand side are defined as:

$V_{\mathrm{maxii}}$ denotes basal glucose utilisation,
$V_{\mathrm{maxid}}$ denotes insulin dependent glucose utilisation, and
$\frac{I}{IS}$ denotes plasma insulin divided by the insulin sensitivity. The division of insulin sensitivity means that if the diseased state is high then the insulin-dependent glucose utilisation is low.

$U_{\mathrm{id}}=\frac{V\max_{U}*\mathrm{Gt}}{IS*U_{50}+ Gt}$	(Eq. S51)
where $U_{\mathrm{id}}$ describes total glucose utilisation from both muscle and fat tissue, and where the components of the right-hand side are defined as:

$V\max_{U}$ denotes maximum glucose utilisation,

$\mathrm{IS}$ denotes impact of disease function,
$U_{50}$ denotes glucose exposure when half of the maximum transportation rate is reached, and
$\mathrm{Gt}$ denotes glucose in tissue.

The total glucose utilisation from both muscle and fat tissue is separated into corresponding muscle and fat glucose utilisation using a static fraction (**Eqs. S52-S53**).

$U_{\mathrm{idm}}=U_{id}*U_{ratio}$	(Eq. S52)
where $U_{\mathrm{idm}}$ describes muscle glucose utilisation, and where the components of the right-hand side are defined as:

$U_{id}$ denotes the total glucose utilisation of both muscle and fat tissue, and
$U_{ratio}$ denotes the fraction of the utilisation which is from the muscle tissue.

$U_{\mathrm{idf}}=U_{id}*(1- U_{ratio})$	(Eq. S53)
where $U_{\mathrm{idf}}$ describes fat glucose utilisation, and where the components of the right-hand side are defined as:
1 represents the inverse of the $U_{ratio}$,

$U_{id}$ denotes the total glucose utilisation of both muscle and fat tissue, and
$U_{ratio}$ denotes the fraction of the utilisation which is from the muscle tissue.

In the model, hepatic glucose can be metabolised into glycogen (**Eq. S54**) and glycogen can be metabolised back into glucose (**Eq. S55**).

$Glycogenesis= \frac{\mathrm{VglyS}_{\max}* Gl}{\mathrm{VglyS}_{50}+Gl}* \mathrm{InsulinResponseNew}$	(Eq. S54)
where $\mathrm{Glycogenesis}$ describes hepatic glycogenesis, and where the components of the right-hand side are defined as:

$\mathrm{VglyS}_{\max}$ denotes maximum rate,
$\mathrm{Gl}$ denotes glucose in liver,
$\mathrm{VglyS}_{50}$ denotes glucose exposure when half of the maximum rate is reached, and I$\mathrm{nsulinResponseNew}$ denotes the new hepatic insulin response.

$Glycogenolysis = \frac{\mathrm{VglyB}_{\max}* Gly_{L}}{\mathrm{VglyB}_{50}+ Gly_{L}}$	(Eq. S55)
where $\mathrm{Glycogenolysis}$ describes hepatic glycogenolysis, and where the components of the right-hand side are defined as:

$\mathrm{VglyB}_{\max}$ denotes maximum rate,
$\mathrm{Gl}y_{L}$ denotes glycogen in liver, and
$\mathrm{VglyB}_{50}$ denotes glucose exposure when half of the maximum rate is reached.

In addition to hepatic glycogen, hepatic glucose can also be metabolised into pyruvate (**Eq. S56)**, and the glucose can indirectly be produced again from gluconeogenesis (**Eq. S57**).

$Glycolysis = Glycolysis_{k}*Gl *InsulinResponseNew$	(Eq. S56)
where $\mathrm{Glycolysis}$ describes hepatic glycolysis, and where the components of the right-hand side are defined as:

$\mathrm{Glycolysi}s_{k}$ denotes basal reaction rate,
$\mathrm{Gl}$ denotes hepatic glucose, and
$\mathrm{InsulinResponseNew}$ denotes the new hepatic insulin response.

$Gluconeogenesis= TCAcycle_{L}*Gluconeogenesis_{\mathrm{TCAk}}*GlyDep_{\mathrm{GluconeogenesisNegative}}$	(Eq. S57)
where $\mathrm{Gluconeogenesis}$ describes hepatic gluconeogenesis, and where the components of the right-hand side are defined as:

$\mathrm{TCAcycl}e_{L}$ denotes a summarised state of components of the TCA cycle,
$\mathrm{Gluconeogenesi}s_{\mathrm{TCAk}}$ denotes basal gluconeogenesis reaction rate, and $\mathrm{GlyDe}p_{\mathrm{GluconeogenesisNegative}}$ denotes a dependency to the fed state of the body. The $\mathrm{GlyDe}p_{\mathrm{GluconeogenesisNegative}}$ is described with a negative dependency to the fed state of the body, which means that gluconeogenesis is upregulated in a fasted state.

Components of the TCA cycle can be metabolised out of the model. This is described in the model with one equation with a positive dependency to the fed state of the body (**Eq. S58**).

$\mathrm{TC}A_{\mathrm{usage}} =TCAcycle_{L}*TCAusage_{K}* GlyDep_{\mathrm{UtilizationPositive}}$	(Eq. S58)
where $\mathrm{TC}A_{\mathrm{usage}}$ describes usage of hepatic TCA cycle components, and where the components of the right-hand side are defined as:
$\mathrm{TCAcycl}e_{L}$ denotes a summarised state of components of the hepatic TCA cycle,
$\mathrm{TCAusag}e_{K}$ denotes the basal reaction rate, and
$\mathrm{GlyDe}p_{\mathrm{UtilizationPositive}}$ denotes a dependency to the fed state of the body. The $\mathrm{GlyDe}p_{\mathrm{UtilizationPositive}}$ is described with a positive dependency to the fed state of the body, which means that gluconeogenesis is downregulated when in a fed state.

Hepatic pyruvate can in the model be metabolised into the TCA cycle (**Eq. S59**) and the hepatic metabolism can be supplied with metabolites that appears in the lactate compartment, which is not sourced from meals (**Eq. S60**).

$\mathrm{PyruvateOx}i_{L}=Pyruvate_{L}* PyruvateOxi_{K}$	(Eq. S59)
where $\mathrm{PyruvateOx}i_{L}$ describes the rate of pyruvate oxidation into the TCA cycle, and where the components of the right-hand side are defined as:
$\mathrm{PyruvateOx}i_{K}$ denotes reaction rate, and
$\mathrm{Pyruvat}e_{L}$ denotes amount of hepatic pyruvate.

$PyruvateTranslocase=PyruvateTranslocase_{K}*GlyDep_{\mathrm{inflow}}$	(Eq. S60)
where $\mathrm{PyruvateTranslocase}$ describes a flow into the pyruvate compartment sourced from the body, and where the components of the right-hand side are defined as:
$\mathrm{PyruvateTranslocas}e_{K}$ denotes basal reaction rate, and
$\mathrm{GlyDe}p_{\mathrm{inflow}}$ denotes a dependency to the fed state of the body. The $\mathrm{GlyDe}p_{\mathrm{inflow}}$ increases the flow into the model during a fasted state. The equation is a simplification of many various reactions in the body to describe the fasting between meals with the least amount of complexity. For example, the breakdown of muscle and fat tissue during long-term fasting is one source that is summarised in the $\mathrm{PyruvateTranslocase}$.

In the model, similarly to the human cell cultures, insulin production is described in three parts: 1) dependency to glucose concentration (**Eq. S61**); 2) dependency to glucose rate of change (**Eq. S62**); and 3) basal insulin production (parameter $S_{b}$). The two glucose-dependent parts of the insulin production are dependent on exenatide exposure, in accordance with the mathematical model used to describe exenatide effect on the human cell cultures.

$ChangeinGlucose=\frac{K}{180.16}*\frac{d}{dt}(Gp)$	(Eq. S61)
where $\mathrm{ChangeinGlucose}$ describes plasma glucose rate of change and where the components of the right-hand side are defined as:
$K$ denotes the insulin production from plasma glucose rate of change,
180.16 denotes glucose molecular weight, and
 $\frac{d}{dt}(Gp)$ denotes plasma glucose rate of change.

As described in this supplementary note above, two alternatives of summarised insulin production equation were tested to describe the glucose injection response in rats. The first alternative (**Eq. S62a**) is the original version of the equation which was used to describe human meal responses, and the proposed alternative was used to describe the glucose injection response in rats (**Eq. S62b**).

$S_{\mathrm{po}}=max(0,ChangeInGlucose*\mathrm{Exenatide}_{\mathrm{IProc}}+{InsulinStabilization*\mathrm{Exenatide}_{\mathrm{IPbasal}}+S}_{b}$)	(Eq. S62a)
where $S_{\mathrm{po}}$ describes the total insulin production, and where the components of the right-hand side are defined as:

$\max$ denotes that function output is always zero or positive,
$\mathrm{ChangeInGlucose}$ denotes the scaled plasma glucose rate of change,
$\mathrm{Exenatide}_{\mathrm{IProc}}$ denotes exenatide effect on glucose rate of change,
$\mathrm{InsulinStabilization}$ denotes the scaled and delayed response of glucose concentration dependent insulin production,
$\mathrm{Exenatide}_{\mathrm{IPbasal}}$ denotes exenatide effect on glucose concentration dependent insulin production, and
$S_{b}$ denotes basal insulin production.

$S_{\mathrm{po}}=\max\left( 0,ChangeInGlucose*\mathrm{Exenatide}_{\mathrm{IProc}} \right)+{max(0,InsulinStabilization*\mathrm{Exenatide}_{\mathrm{IPbasal}})+S}_{b}$	(Eq. S62b)

where all the variables are described in the original equation (**Eq. S62a**). The proposed alternative includes a minimalistic change which provided a greater agreement to rat glucose injection data. This difference in agreement to data may be because the original equation was not developed to describe glucose injections, where glucose rate of change is much higher than meal responses. The high glucose rate of change may disturb the insulin production balance between the three parts of the insulin production equations, which could be the first alternative had a lower ability to describe the data. As described in this supplementary note above, in practice, the alternative equation means that negative outputs of either $\mathrm{ChangeInGlucose}$ or $\mathrm{InsulinStabilization}$ cannot reduce the overall insulin production – which was favourable when fitting the model to the rat glucose injection study. The testing of both versions of the model alternatives are included when downloading the model from our GitHub repository.

In the model, insulin secretion is described with one equation (**Eq. S63**).

$S =gamma*Ipo$	(Eq. S63)
where $S$ describes insulin secretion, and where the components of the right-hand side are defined as:
$\mathrm{gamma}$ denotes transportation rate, and
$\mathrm{Ipo}$ denotes insulin in the portal vein.

Insulin transportation between the liver and plasma is described by two equations (**Eq. S64-S65**).

$IpToIl =m_{2}*Ip$	(Eq. S64)
where $\mathrm{IpToIl}$ describes insulin rate from plasma to the liver, and where the components of the right-hand side are defined as:
$m_{2}$ denotes transportation rate, and
$\mathrm{Ip}$ denotes insulin in the plasma.

$IlToIp=m_{1}*Il$	(Eq. S65)
where $\mathrm{IlToIp}$ describes insulin rate from liver to plasma, and where the components of the right-hand side are defined as:
$m_{1}$ denotes transportation rate, and
$\mathrm{Il}$ denotes insulin in the liver.

Insulin clearance is described as a sum of hepatic clearance (**Eq. S66-S68**) and non-hepatic clearance (**Eq. S69**). The hepatic and non-hepatic clearance is sourced from the original model and for a deeper description of the four equations we refer to the source of the equations (15).

$HE =max(0,-m_{5}*S+m_{6})$	(Eq. S66)
where $\mathrm{HE}$ describes the dynamic part of the hepatic excretion, and where the components of the right-hand side are defined as:
$\max$ denotes that the output is always zero or positive,
$m_{5}$ denotes hepatic clearance reduces as insulin secretion increases,
$S$ denotes insulin secretion, and
$m_{6}$ denotes basal reaction rate.

$M_{3} =\frac{HE *m_{1}}{1 - HE}$	(Eq. S67)
where $M_{3}$ is the reaction rate of hepatic excretion, and where the components of the right-hand side are defined as:
$\mathrm{HE}$ denotes the dynamic part of the hepatic excretion, and
$m_{1}$ denotes the maximum hepatic excretion rate.

$\mathrm{InsulinDegradation}_{\mathrm{Liver}}=M_{3}*\mathrm{Insuli}n_{\mathrm{Liver}}$	(Eq. S68)
where $\mathrm{InsulinDegradation}_{\mathrm{Liver}}$ describes the hepatic insulin clearance, and where the components of the right-hand side are defined as:
$M_{3}$ denotes the reaction rate, and
$\mathrm{Insuli}n_{\mathrm{Liver}}$ denotes insulin in the liver.

$\mathrm{InsulinDegradation}_{\mathrm{Blood}}=m_{4}*\mathrm{Ip}$	(Eq. S69)
where $\mathrm{InsulinDegradation}_{\mathrm{Blood}}$ describes non-hepatic insulin clearance, and where the components of the right-hand side are defined as:$m_{4}$ denotes reaction rate, and
$\mathrm{Ip}$ denotes insulin in plasma.

The final component of the model is model variables.

##### e) Model variables

The four metabolic variables (two volumes and two metabolic rates) which was to scaled in the pharmacokinetic model was implemented in the M4-model to identically scale exenatide pharmacokinetics to total body weight (**Eqs. S7-S10**).

Similarly to the pharmacokinetic model, blood volumes was also scaled to total body weight (**Eqs. S70-71**).

$PlasmaVolume =$	$0.02176* {(\frac{BW}{0.34})}^{1.06}$	(Eq. S70)

where $\mathrm{PlasmaVolume}$ describes plasma volume, and where the components of the right-hand side are defined as:

$0.02176$ denotes assumed rat total blood volume (L) as a reference species with a body weight assumed to be 0.34 kg (17,18),
$BW$ denotes subject total body weight, and
$1.06$ denotes allometric scaling constant. The allometric scaling constant of 1.06 was set in order for the model equation output to be identical to the original model equation at 80 kg. Note that this part of the model is a larger simplification of the metabolism and was only set to generate the same volume output as the original model. The assumption has not been tested to describe body weights between 0.34 kg (rat study) and 102 kg (human study), and therefore we recommend setting the plasma volume manually to your own estimates, if using the model for simulations not plotted in the manuscript.

$LiverVolume=PlasmaVolume*0.13$	(Eq. S71)
where $\mathrm{LiverVolume}$ describes blood volume in the liver, and where the components of the right-hand side are defined as:
$0.13$ denotes 13% of the plasma blood volume is in the liver, and
$\mathrm{PlasmaVolume}$ denotes plasma volume. Note that this equation is a simplification, where not all animal species or humans have the same relation between liver and plasma blood volumes. This simplification was unchanged from the pre-existing model (4), and was left unchanged to minimise changes to the model. We recommend setting the liver volume manually to your own estimates, if using the model for simulations not plotted in the manuscript.

Hepatic glycogen concentrations are estimated with a unit conversion from the state $\mathrm{Gl}y_{L}$ (**Eq. S72**).

$\mathrm{Glycogen}_{\mathrm{Liver}} =\frac{\mathrm{Gl}y_{L}*\mathrm{BW}}{\mathrm{VolumeBloo}d_{\mathrm{Liver}} * \mathrm{MolecularWeight}_{\mathrm{Glycogen}}}$	(Eq. S72)

where $\mathrm{Glycogen}_{\mathrm{Liver}}$ describes hepatic glycogen concentration (mM), and where the components of the right-hand side are defined as:

$\mathrm{Gl}y_{L}$ denotes hepatic glycogen (mg/kg),
$\mathrm{BW}$ denotes total species body weight,
$\mathrm{LiverVolume}$ denotes liver volume, and
$\mathrm{MolecularWeight}_{\mathrm{Glycogen}}$ denotes molecular weight of glycogen which is assumed to be 666.6 Da. Note that glycogen is assumed to have a molecular weight of 666.6 which is a simplification which originates from the pre-existing model (4).

Plasma insulin concentration is estimated with a unit conversion from the state $\mathrm{Ip}$ (**Eq. S73**).

$\mathrm{Insuli}n_{\mathrm{Blood}}=\frac{Ip* BW}{\mathrm{PlasmaVolume}}$	(Eq. S73)

where $\mathrm{Insuli}n_{\mathrm{Blood}}$describes plasma insulin concentration (pM), and where the components of the right-hand side are defined as:
$\mathrm{Ip}$ denotes insulin in the plasma (pmol/kg),
$\mathrm{BW}$ denotes subject total body weight, and
$\mathrm{PlasmaVolume}$ denotes total blood volume.

Plasma glucose concentration is estimated with a unit conversion from the state $\mathrm{Gp}$ (**Eq. S74**).

$\mathrm{Glucos}e_{\mathrm{Blood}} =\frac{Gp * BW}{\mathrm{PlasmaVolume} *10}$	(Eq. S74)
where $\mathrm{Glucos}e_{\mathrm{Blood}}$ describes the typically American-used plasma glucose unit (mg/dl), and where the components of the right-hand side are defined as:

$\mathrm{Gp}$ denotes glucose in the plasma (mg/kg),
$\mathrm{BW}$ denotes subject total body weight, and
$PlasmaVolume*10$ denotes assumed total blood volume (dl). The standardised American measurement (mg/dl) was converted into the standardised European glucose measurement (mM) through a division of 18.

One final remark of the model is the various regulations in the model based on the fed state of the body. These regulations are simplifications which is unchanged from the original model (**Eqs. S75-79**) (4).

${\mathrm{GlyDe}p_{\mathrm{MealPositive}}=\left( \frac{\mathrm{Glycogen}_{\mathrm{Liver}}}{\mathrm{GlyDe}p_{\mathrm{Meal}}} \right)}^{\mathrm{GlyDepEXP}_{\mathrm{Meal}}}$	(Eq. S75)
where $\mathrm{GlyDe}p_{\mathrm{MealPositive}}$ describes a positive dependency to the fed state of the body, and where the components of the right-hand side are defined as:

$\mathrm{Glycogen}_{\mathrm{Liver}}$ denotes hepatic glycogen,
$\mathrm{GlyDe}p_{\mathrm{Meal}}$ denotes a hepatic glycogen baseline where the function will not affect the reactions dependent on $\mathrm{GlyDe}p_{\mathrm{MealPositive}},$and
$\mathrm{GlyDepEXP}_{\mathrm{Meal}}$ denotes scaling between the hepatic glycogen levels and the function output. $\mathrm{GlyDe}p_{\mathrm{MealPositive}}$ was used to upregulate flow of amino acids away from hepatic gluconeogenesis when consuming protein in a fed state.

$\mathrm{GlyDe}p_{\mathrm{MealNegative}}= \left( \frac{\mathrm{GlyDep}_{\mathrm{Meal}}}{\mathrm{Glycogen}_{\mathrm{Liver}}} \right)^{\mathrm{GlyDepEXP}_{\mathrm{Meal}}}$	(Eq. S76)
where $\mathrm{GlyDe}p_{\mathrm{MealNegative}}$ describes a negative dependency to the fed state of the body, and where the components of the right-hand side are defined as:

$\mathrm{Glycogen}_{\mathrm{Liver}}$denotes hepatic glycogen,
$\mathrm{GlyDe}p_{\mathrm{Meal}}$ denotes a hepatic glycogen baseline where the function will not affect the reactions dependent on $\mathrm{GlyDe}p_{\mathrm{MealNegative}},$and
$\mathrm{GlyDepEXP}_{\mathrm{Meal}}$ denotes scaling between the hepatic glycogen levels and the function output. $\mathrm{GlyDe}p_{\mathrm{MealNegative}}$ was used to upregulate flow of amino acids into the gluconeogenesis when consuming protein in a fed state.

${\mathrm{GlyDe}p_{\mathrm{UtilizationPositive}} =\left( \frac{\mathrm{Glycogen}_{\mathrm{Liver}}}{\mathrm{GlyDe}p_{\mathrm{TCA}}} \right)}^{\mathrm{GlyDepEXP}_{\mathrm{TCA}}}$	(Eq. S77)
where $\mathrm{GlyDe}p_{\mathrm{UtilizationPositive}}$ describes a positive dependency to the fed state of the body, and where the components of the right-hand side are defined as:

$\mathrm{Glycogen}_{\mathrm{Liver}}$ denotes hepatic glycogen,
$\mathrm{GlyDe}p_{\mathrm{Utilization}}$ denotes a hepatic glycogen baseline where the function will not affect the reactions dependent on $\mathrm{GlyDe}p_{\mathrm{MealPositive}},$and
$\mathrm{GlyDepEXP}_{\mathrm{Utilization}}$ denotes scaling between the hepatic glycogen levels and the function output. $\mathrm{GlyDe}p_{\mathrm{UtilizationPositive}}$ was used to upregulate the usage of components in the hepatic TCA cycle in a fed state.

$\mathrm{GlyDe}p_{\mathrm{GluconeogenesisNegative}}= \left( \frac{\mathrm{GlyDe}p_{\mathrm{Gluconeogenesis}}}{\mathrm{Glycogen}_{\mathrm{Liver}}} \right)^{\mathrm{GlyDepEX}P_{\mathrm{Gluconeogenesis}}}$	(Eq. S78)
where $\mathrm{GlyDe}p_{\mathrm{GluconeogenesisNegative}}$ describes a negative dependency to the fed state of the body, and where the components of the right-hand side are defined as:

$\mathrm{Glycogen}_{\mathrm{Liver}}$ denotes hepatic glycogen,
$\mathrm{GlyDe}p_{\mathrm{Gluconeogenesis}}$ denotes a hepatic glycogen baseline where the function will not affect the reactions dependent on $\mathrm{GlyDe}p_{\mathrm{GluconeogenesisNegative}},$and
G$\mathrm{lyDepEX}P_{\mathrm{Gluconeogenesis}}$ denotes scaling between the hepatic glycogen levels and the function output. $\mathrm{GlyDe}p_{\mathrm{GluconeogenesisNegative}}$ was used to upregulate the renal glucose production in a fed state.

$\mathrm{GlyDe}p_{\mathrm{inflow}}=\left( \frac{\mathrm{GlyDe}p_{\mathrm{KidneysEGP}}}{\mathrm{Glycogen}_{\mathrm{Liver}}} \right)^{\mathrm{GlyDepEX}P_{\mathrm{KidneysEGP}}}$	(Eq. S79)
where $\mathrm{GlyDe}p_{\mathrm{inflow}}$ describes a negative dependency to the fed state of the body, and where the components of the right-hand side are defined as:

$\mathrm{Glycogen}_{\mathrm{Liver}}$ denotes hepatic glycogen,
$\mathrm{GlyDe}\mathrm{pIn}_{K}$ denotes a hepatic glycogen baseline where the function will not affect the reactions dependent on $\mathrm{GlyDe}p_{\mathrm{inflow}},$and
$\mathrm{GlyDepI}n_{\mathrm{kEGP}}$ denotes scaling between the hepatic glycogen levels and the function output. $\mathrm{GlyDe}p_{\mathrm{inflow}}$ was used to both downregulate the renal glucose production and the flow of components into the hepatic pyruvate compartment in a fed state.

Together, all these described equations (Eqs. S1–S79) constitute the full M4-model used in the main text for our analysis.
